# Dynamic simulations of transcriptional control during cell reprogramming reveal spatial chromatin caging

**DOI:** 10.1101/642009

**Authors:** Marco Di Stefano, Ralph Stadhouders, Irene Farabella, David Castillo, François Serra, Thomas Graf, Marc A. Marti-Renom

## Abstract

Chromosome structure is a crucial regulatory factor for a wide range of nuclear processes. Chromosome Conformation Capture (3C)-based experiments combined with computational modelling are pivotal for unveiling 3D chromosome structure. Here, we introduce TADdyn, a new tool that integrates time-course 3C data, restraint-based modelling, and molecular dynamics to simulate the structural rearrangements of genomic loci in a completely data-driven way. We applied TADdyn on *in-situ* Hi-C time-course experiments studying the reprogramming of murine B cells to pluripotent cells, and characterized the structural rearrangements that take place upon changes in the transcriptional state of 11 genomic loci. TADdyn simulations show that structural *cages* form around the transcription starting site of active loci to stabilize their dynamics, by initiating (*hit*) and maintaining (*stick*) interactions with regulatory regions. Consistent findings with TADdyn for all loci under study suggest that this *hit-and-stick* mechanism may represent a general mechanism to trigger and stabilize transcription.

## INTRODUCTION

The three-dimensional (3D) structure of the genome has been shown to modulate transcriptional regulation^1–3^ and to play a role in cancer and developmental abnormalities^4^. In the effort of characterising 3D genome structures, Chromosome Conformation Capture (3C)-based experiments^5^ allow to capture a single snapshot of the genome conformation at a given time. A plethora of theoretical approaches have been developed to take advantage of 3C-based experimental data and model genome spatial organization. Restraint-based modelling approaches^6^ take 3C-based contact frequencies as input and employ *ad hoc* conversions to spatial distances for determining 3D genome structure^7–12^. This approach has provided valuable insights into the structural organization of chromosomal regions in various organisms^13^. Complementary, thermodynamics-based approaches^14–22^ use physics-based principles to test specific interactions or interaction mechanisms to explain the molecular origins of the contact patterns obtained in 3C-based experiments. Together, these theoretical approaches provide insights into chromatin conformation^16,17,23,24^ and the possible mechanisms that form chromosome territories^18^, compartments^19^ and Topologically Associating Domains (TADs)^20,22,25,26^.

Decreased sequencing costs together with more refined experimental protocols has recently permitted performing 3C-based time-resolved experiments to monitor genome conformation dynamics of biological processes at high resolution. For example, Hi-C experiments have been applied to study nuclear organization during mitosis^27,28^ or meiosis^29–31^, perturbations induced by hormone treatment^32^ and during induced neural differentiation^33^ or cell reprogramming^34^. However, none of the strategies mentioned above can currently take full advantage of these new datasets, and computational approaches specifically designed for the simulation of time-dependent conformational changes (4D) are not yet available. To fill this gap, we introduce TADdyn, a novel computational method which allows modeling 3D structural transitions of chromatin using time-resolved Hi-C datasets combining restraint-based modelling and molecular dynamics. We found two distinct phases of 3D genome conformation dynamics by applying TADdyn to 11 representative gene loci during cell reprogramming of mouse pre-B lymphocytes into Pluripotent Stem Cells (PSCs)^34^. First, the transcription start site (TSS) forms a ‘cage’-like structure within a TAD, favoring specific contacts with open and active (enhancer) regions that may be located several kilo-bases (kb) away (here called the ‘hit phase’). Second, the cage confines and stabilizes the dynamics of the TSS and its interactions with active regions against simulated thermal motion (the ‘stick phase’). We propose this hit-and-stick model as a general mechanism for chromatin architectural changes linked to transcriptional activation.

## RESULTS

### The TADdyn modelling strategy

TADdyn is based on the following methodological steps (**Methods** and **Fig. 1**): (i) collection of experimental data, (ii) representation of selected chromatin regions using a bead-spring polymer model, (iii) conversion of experimental data into time-dependent restraints, (iv) application of steered molecular dynamics to simulate the adaptation of chromatin models to satisfy the imposed restraints, and (v) analysis of the spatial conformations obtained.

**Figure 1.**
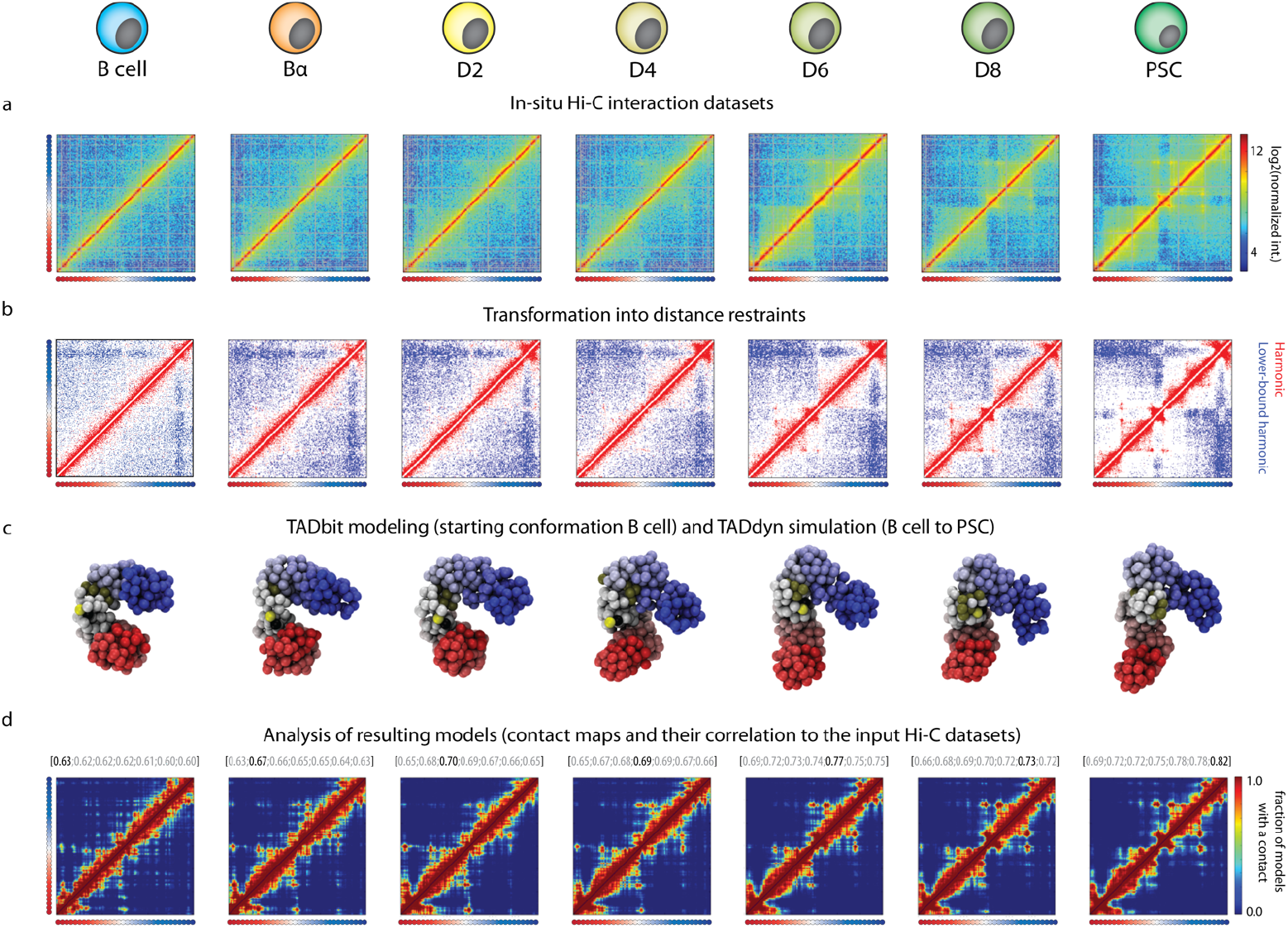
The TADdyn key steps for simulating 4D dynamic changes in a *locus*. The shown example corresponds to the reprogramming dynamics for the *Sox2* locus from B cells to PSC. **a** Data collection. *In-situ* normalized Hi-C interaction matrices34 for the region of 1.5Mb centered around the *Sox2* promoter. **b** Distance restraints definition. Both *LowerBound*- (blue) and *Harmonic* (red) spatial restraints are obtained by filtering the Hi-C interaction maps using the optimal triplet of TADbit parameters. **c** TADdyn steered dynamics runs. The *loci* are represented as polymers made of spherical particles containing 5kb each a bin of the input Hi-C matrix. Particles are colored from red (start of the modeled region) to blue (end of the modeled region). The TSS of the locus of interest is represented by a black particle. The models for all the stages (that is, from B to PSC) were dynamically built by TADdyn. Only the models at the experimental time-points are shown, but TADdyn allows to visualize the entire dynamics *filling the blanks* between stages (**Suppl. Videos 1-11**). **d** Contact maps (<200 nm distance cutoff) of models during the simulation were used to assess their accuracy by means of its correlation with the Hi-C input matrices. The Spearman correlation coefficients (SCCs) of the contact maps with the seven Hi-C input matrices are shown with the coefficient in black letters corresponding to the time point of the column.

We applied TADdyn to a previously published *in-situ* Hi-C interaction time-series dataset (GEO accession number GSE96611). This included seven time-points of *in-situ* Hi-C experiments during C/EBPα priming followed by Oct4, Sox2, Klf4 and Myc (OSKM)-induced reprogramming of B cells to PSCs^34^. We focused on 10 different ~2Mb regions of the mouse genome encompassing the promoters of 11 genes of interest. We selected these specific genes because they are representative of different time-dependent patterns of transcriptional activity, which allowed us to study how various different transcription dynamics interplay with changes in the 3D genomic organization. Specifically, we analyzed the early activated *Tet2* gene; the late activated *Sox2* and *Nanog* genes; the transiently activated *Mmp3*, *Mmp12*, *CEBPα* and *Nos1ap* genes; the transiently silenced *Lmo7* genes, the stably silenced *Ebf1* gene and the essentially invariant genes *Rad23a* and *Rad23b* (**Suppl. Figs. 1-11a** and **Table 1**).

Next, experimental Hi-C interaction matrices of the 10 regions at 5kb resolution were converted into TADdyn time-dependent spatial restraints (**Fig. 1a**). Specifically, each 5kb-bin was represented as a spherical particle of diameter 50nm, and at each experimental stage the Hi-C interaction matrix was converted into harmonic spatial restraints between these particles (**Fig. 1b**). This conversion follows the simple, yet effective, rationale that pair of particles with a high number of Hi-C interactions between them are restrained to stay close in space, while poorly contacting particle pairs are kept far apart^8^ (**Methods**). The parameters of each imposed restraint between two particles (that is, the spring constant and its equilibrium distance) are then linearly interpolated between the values of the consecutive experimental time points using steered molecular dynamics simulations. This strategy favors the adaptation of the chromatin models to the imposed dynamic restraints (**Methods** and **Suppl. Fig. 12**). TADdyn restraints are specifically designed to allow smooth structural changes between models of consecutive experimental time-points.

### TADdyn simulations are an accurate dynamic representation of the input Hi-C data

To quantify the degree of agreement of the TADdyn 4D models to the time-series Hi-C interaction matrices, contact maps were calculated on an ensemble of 100 models obtained at each time step of the trajectories (**Fig. 1d**). Next, for each time-point the Spearman correlation coefficient (SCC) of the modeled contact map and the experimental Hi-C interaction matrices was computed as a measure of the agreement between the two interactions matrices (**Methods**). For the 11 loci, the SCC ranged between 0.60 and 0.89, which is indicative of accurate models^35^ (**Fig. 1d** for *Sox2* simulations and **Suppl. Fig. 1-11c** for all other loci). Notably, at a fixed cell stage, the Hi-C interaction map correlated best with the contacts maps at the corresponding time of the simulated reprogramming dynamics. These results show that for a diverse set of loci the simulated structures are a reliable 3D representation of two-dimensional Hi-C interaction matrices, which effectively reproduce the actual contact pattern at the correct time of the trajectory.

To test whether TADdyn is suited to model processes characterized by gradual chromatin structural transitions, two alternative sets of 100 simulations each for *Sox2* were performed by completely deleting the restraints of stages D2 (ΔD2 simulation) and D6 (ΔD6 simulation). By doubling the time duration between the remaining adjacent transitions (Bα to D4 and D4 to D8), the total duration of the trajectories was maintained constant. We observed that the missing restraints only marginally affected the results of the simulations. Specifically, the conformations expected to represent the missing cell stages along the trajectories still provided accurate models at the removed stage (SCC_D2_=0.68 and SCC_D6_=0.75 in ΔD2 and ΔD6 simulations, respectively). Additionally, in the case of ΔD2, the Hi-C interactions map at D2 correlated best with the contact map computed on the models predicting the D2 stage (for ΔD6 the SCC between Hi-C at D6 and models in ES correlated slightly better than the one for D6; SCC_ES_=0.76 *vs.* SCC_D6_=0.75). These results suggest that the simulated reprogramming dynamics are dominated by smooth and gradual chromatin rearrangements that can be effectively simulated by TADdyn.

In the following sections, we present the simulation details the simulations for three representative loci: *Sox2* (as a late activated *locus*), *Mmp12* (as a late repressed locus) and *Rad23a* (as a stably active *locus*). The detailed results for all the simulated loci are presented in the **Suppl. Figs 1-11** and summarized in the last paragraph of the **Results** section.

### Dynamic structural reorganization correlates with local transcriptional changes

To explore how the simulated models changed over time, we performed a hierarchical clustering analysis of the correlation matrix between the contact maps of the models and the input Hi-C interaction matrices (**Methods**, **Fig. 2a** and **Suppl. Figures 1-11d**). Hi-C interaction matrices were grouped in well-separated clusters reflecting the expected changes in expression activity of each locus. This clustering indicates that the studied loci changed their topology along with their transcriptional activity. Specifically, for 8 of the 11 simulated loci (that is *CEBPα*, *Ebf1*, *Lmo7*, *Mmp12*, *Mmp3*, *Nos1ap*, *Sox2*, and *Tet2*) the active and inactive states of the locus were represented by two major clusters. For example, the first cluster of *Sox2* comprised the cell stages from B to D4 in which the locus is inactive (RPKM<0.06), while the second cluster includes D6 to PSC when *Sox2* is active (RPKM>8.0) (**Fig. 2a**). Similar observations were made for *Mmp12*. In contrast, *Rad23a* showed a less clear partition into two distinct clusters (the 3 first clusters are almost equidistant), likely because the locus remains in an active state during reprogramming with relatively small fluctuations of expression. To further characterize the structural changes associated with changes in transcriptional activity, we performed a clustering analysis of the model structures based on the distance RMSD (dRMSD) values between pairs of 3D models (**Methods)**. As for the matrix-based analysis, the dRMSD clustering reflects the presence of different folding states that correlate with transcriptional activity (**Fig. 2b** and **Suppl. Figures 1-11e**).

**Figure 2.**
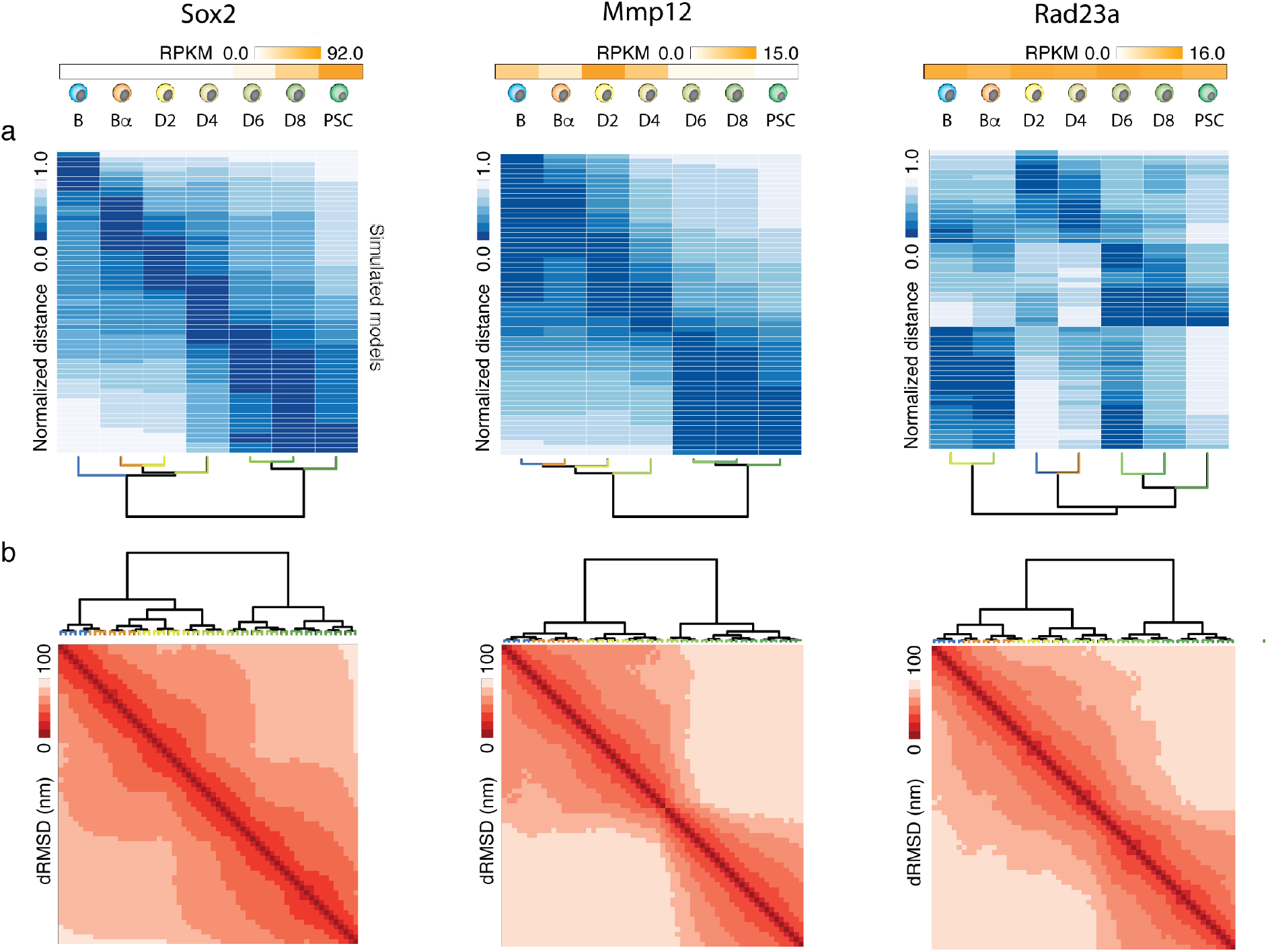
Dynamic structural chromatin reorganization linked to transcriptional activity. Top of the figure indicates expression level (RPKM) for each of the selected loci at each reprogramming stage^34^. **a.** Normalized spearman rank correlation profiles computed for each pair of 60 model contact maps, that is one map every 1 time step of the simulation, against each of the seven Hi-C interaction maps obtained during the reprogramming process. **b.** Model 3D structural clustering based on distance RMSD (dRMSD) between all models within a simulation.

### Gene activation correlates with the appearance of “*cage”*-like structures

In our simulations (**Supplementary videos 1-11**), we observed a *“caging”* effect of the transcription start site (TSS) at the time when a given locus was transcriptionally active. To quantify this observation, we first calculated for each TSS the time-dependent changes in its 3D structural embedding as well its explored volume (**Methods**, **Fig. 3** and **Suppl. Figs. 1-11f,g**). Notably, TADdyn simulations showed that, as cells reprogramed, the TSS for all 11 loci remained largely accessible to other particles during their less transcriptionally active phases (*e.g.* stages B-D4 for *Sox2* and D6-PSC for *Mmp12*) (**Fig. 3a**). During the most active stages the TSS particle became embedded in the 3D model (the TSS embedding was between 0.8-0.95 at the PSC stage for *Sox2* and B-D4 for *Mmp12*) (**Fig. 3a**). In contrast, the embedding profile of the *Rad23a* locus was maintained around 0.6 during the entire simulation in parallel with the overall constant transcriptional activity of the gene.

**Figure 3.**
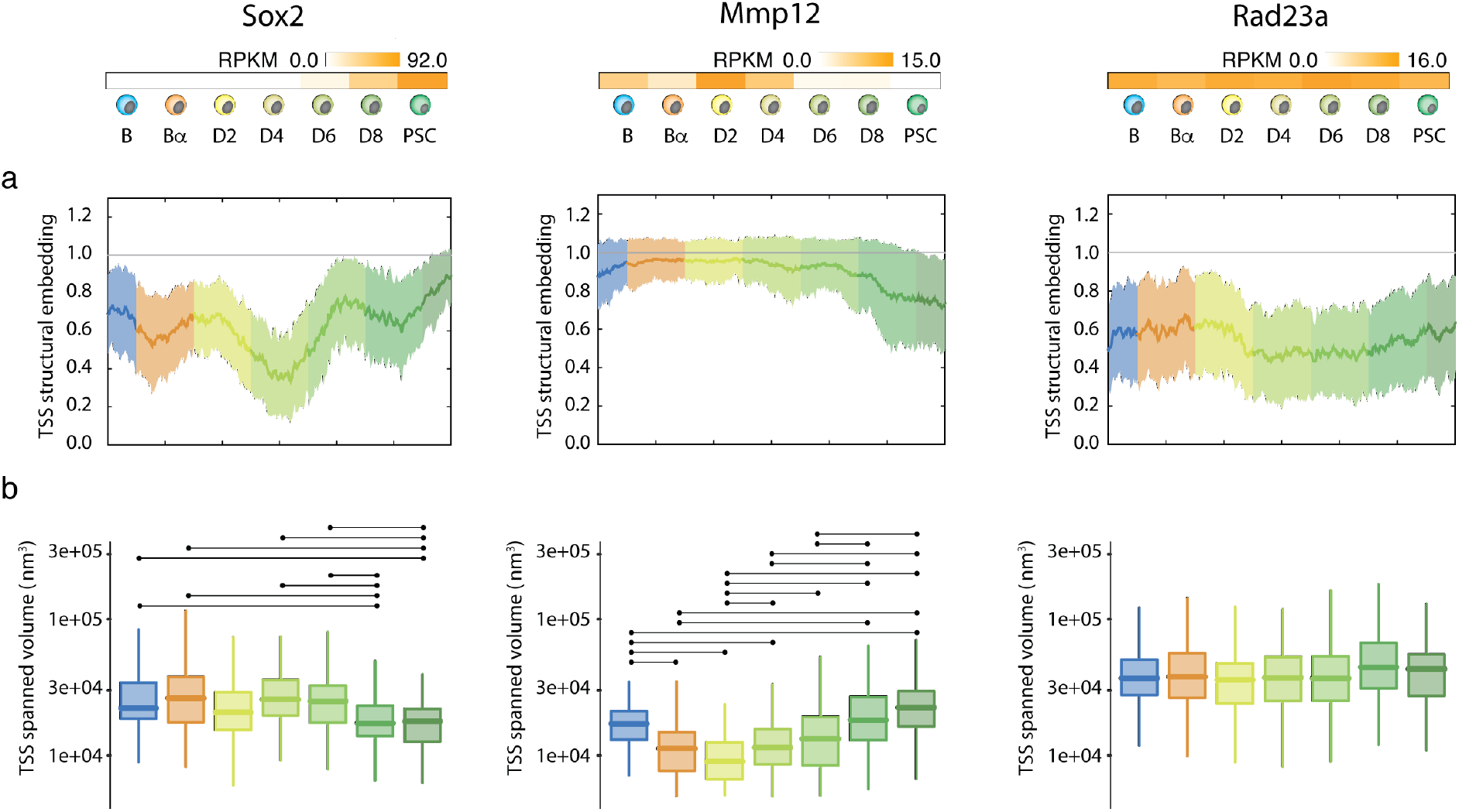
Gene activity correlates with TSS structural embedding and spanned volume. Top of the figure indicates expression level (RPKM) for each of the selected loci at each reprogramming stage^34^. **a** Dynamic structural embedding of the TSS particle. **b** Volume explored by the TSS particle during the various stages of the reprogramming dynamics. Connecting lines between distributions indicate they are statistically different (*p*<0.01, Wilcoxon rank sum test, **Methods**).

Next, we assessed whether the 3D embedding would also affect the spatial dynamics of the TSS from different loci. For each simulation, we calculated the convex-hull of the TSS particle every 50 time-steps of the trajectory as a proxy of the volume explored by the TSS at each reprogramming stage (**Methods** and **Fig. 3b**). The results indicate that the TSS of *Sox2* explores a smaller volume in the D6-PSC stages, when the gene is transcriptionally active, compared to the volume explored in the B to D4 stages when the gene is transcriptionally silent. The TSS of *Mmp12* acquired increased mobility during reprogramming as its expression levels decreased (compare B-D4 and D6-PSC stages). Interestingly, the explored volume of the *Rad23a* TSS barely changed during the entire reprograming process. Taken together, these findings, are consistent with previous super-resolution imaging works suggesting that the mobility of the promoter is constrained upon activation^36,37^.

### Gene activity correlates with domain borders dynamics and enhancer proximity

To further characterize the topological transitions between gene expression states, we calculated dynamic changes in TAD insulation score and border strength^38^ using the contact maps of the models along the TADdyn trajectories (**Methods**). We found that although the number of borders was very constant during the simulation their position often changed. In a striking example involving the *Sox2* locus, during the D4-D6 transition there was a clear shift of a border at position chr3:34.60Mb, moving further downstream and converging near the TSS (**Fig. 4a**). A second border at position chr3:34.74Mb was displaced about 20Kb downstream, thereby inducing the topological insulation of a super-enhancer region at chr3:34.70Mb and the TSS (**Fig. 4b**). These border changes resulted in the formation of a domain of about 120kb, which included exclusively the *Sox2* gene and its super-enhancer region. The fact that the border strength remained high after gene activation at D6 is in agreement with our previously described findings that the *Sox2* TSS and its super-enhancer became isolated in a sub-TAD when the gene is transcribed in PSCs (**Fig. 4a**). Consistent with this observation, the *Mmp12* TSS was partially included in a weak domain border at chr9:7.25Mb during transcription. This weak border disappeared during the gene’s transition from an active to an inactive state (during the D4-D6 stages), so that the *Mmp12* TSS became part of a larger domain at the time of gene silencing. Finally, and with a similar trend, the TSS of the invariantly active gene *Rad23a* remained part of a domain border, and insulated in an unvaried domain between chr8:84.58-82.86Mb for the entire simulation.

**Figure 4.**
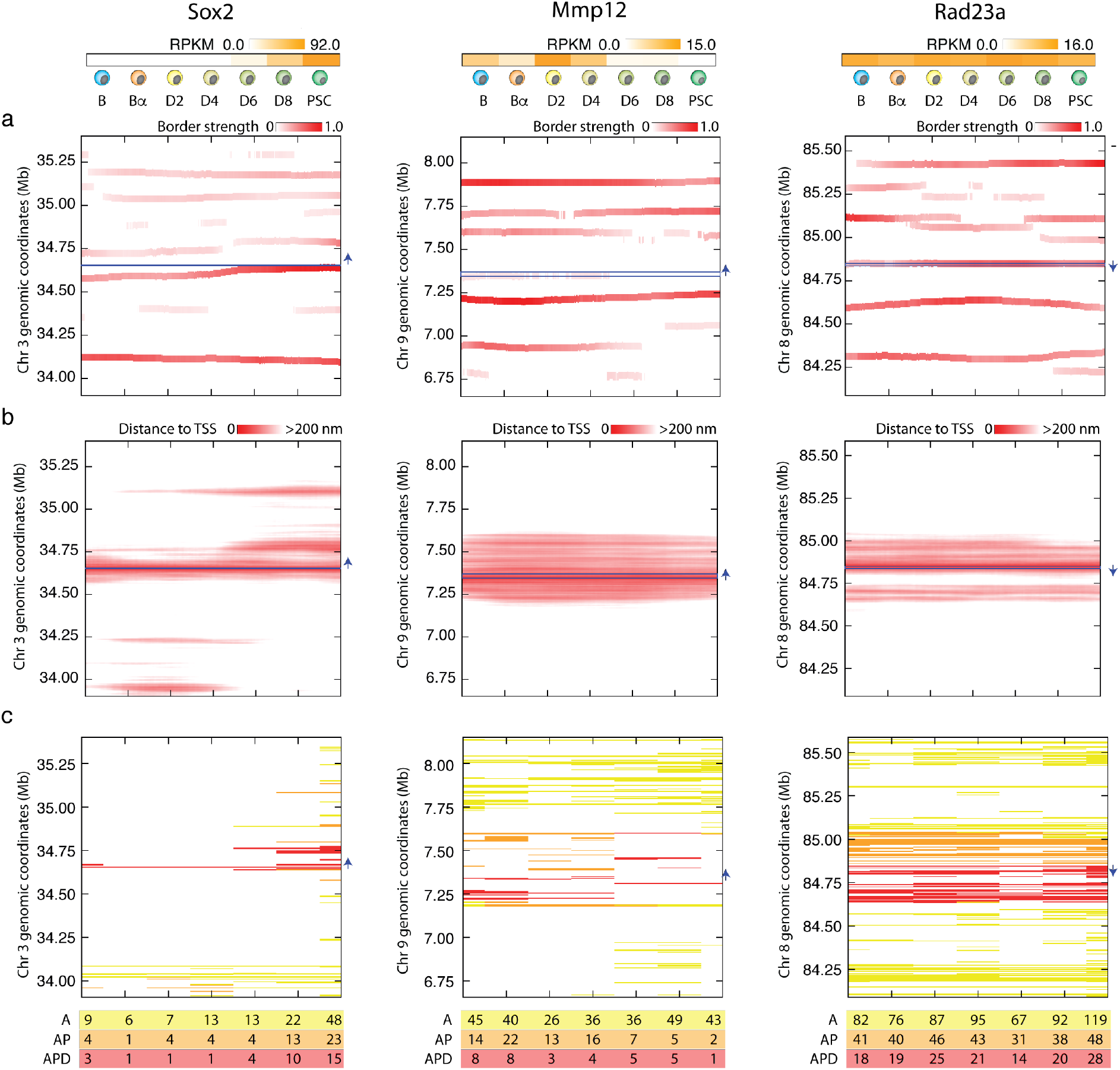
Gene activity correlates with domain borders and enhancer proximity. Top of the figure indicates expression level (RPKM) for each of the selected loci at each reprogramming stage^34^. **a** Time dependent position of the domain borders as defined by the insulation score analysis on the contact maps derived from the models. The position of the loci in the graph is indicated by a blue horizontal line and its transcriptional orientation is indicated by a blue arrow. **b** Heat maps of the distances between the TSS particle and all the other model particles as a function of time. **c** *Active* particles classification into proximal to the TSS (AP, yellow), within the domain (AD, orange), and within the domain and proximal to the TSS (APD, red). The bottom table shows the number of particles in each category for each reprogramming stage.

To characterize the elements within the borders of TADs containing transcriptionally active loci, we first identified in the models what we call “*active*” particles, which had overlapping signals (at least 250bp) of ATAC-seq and H3K4me2 ChIP-seq in any time-point of the reprogramming process^34^ (**Methods**, **Fig. 4b** and **Suppl. Figures 1-11g**). Upon activation, the *Sox2* gene was positioned spatially close (<200nm) to a set of particles containing its super-enhancer (SE) about 120kb downstream of the TSS. Notably, and in line with our previous observations^34^, the TSS-SE proximity started at D4, that is before the *Sox2* transcriptional activity could be detected (D6). This spatial proximity was maintained for the rest of the simulation while the *Sox2* gene was transcribed. In contrast, the TSS 3D distance profiles of *Rad23a* and *Mmp12* remained overall constant during reprogramming (**Fig. 4b**) with marginal changes during the simulation.

To further assess to what extend *Active* particles (**Table 2**) became proximal to the TSS of the locus, we counted for each reprogramming stage how many of the *Active* (A) particles became available to the TSS particle, either by spatial proximity only (*Active-Proximal* particles, AP) or also within the same local domain (*Active-Proximal-Domain*, APD) (**Methods** and **Fig. 4c**). This analysis resulted in a robust trend for all analyzed *loci*, showing higher numbers of AP and APD particles when the *locus* is transcriptionally active. Notably, for *Sox2* the A, AP and APD particles were increasing together with the locus transcription activity, but the regions containing the annotated super-enhancer were only classified as AP or APD at the D6 stage, that is after the structural changes observed during the simulation at D4 stage. For the *Mmp12* simulation the number of A particles did not decrease in the inactive stage, yet the numbers of AP or APD decreased consistently with the general trend. The Rad23a locus was instead characterized by a quite constant and high number of A, AP and APD particles consistently with is stable transcription activity during the entire reprogramming process. Altogether, we observed a consistent 3D co-localization of *Active* and TSS chromatin particles upon activation of the gene, reminiscent of the recently proposed formation of enhancer hubs or condensates^39–41^.

### TADdyn simulations indicate that TSS *caging* is a general structural signature of transcription activation

To move from anecdotal observations to more general trends, we compared relative expression levels with the dynamic chromatin conformation changes detected during the simulations of all 11 selected loci (**Methods** and **Fig. 5**). More specifically, we rescaled RPKM gene expression as well as the other four features obtained from the TADdyn models (that is, the 3D embedding of the TSS particle, the distance to TSS, the volume explored by the TSS, and the number of APD particles) to a 0-to-1 relative scale. Each feature was analyzed for two categories of loci: those that have changes in expression and those that remain active during the entire simulation. The *switch* category includes the loci that show a substantial change in expression level from high values (RPKM>1.0) to low ones (RPKM<0.5) or vice versa, and include *CEPBα*, *Mmp3*, *Mmp12*, *Nos1ap*, *Ebf1*, *Sox2*, and *Nanog*. The *active* simulations were constituted by loci whose activity is always above 1.0 RPKM and included *Lmp7*, *Rad23a*, *Rad23b*, and *Tet2*. Analysis of these 11 *loci* revealed that TSS particles tend to be embedded inside the model structure and closer to other active particles when the gene’s expression is higher (**Fig. 5a, b**), exploring less volume (**Fig. 5c**) and interact with greater numbers of surrounding APD particles (**Fig. 5d**). These findings suggest that the signatures of the *caging* effect are common to all 11 *loci* studied. Additionally, the finding that the numbers of both the AP and APD particles increased with elevated transcription activity (**Figs. 4c** and **5d**) indicates that the formation of a structural cage may have a functional role in promoting and maintaining interactions of the TSS with regulatory regions of the genome.

**Figure 5.**
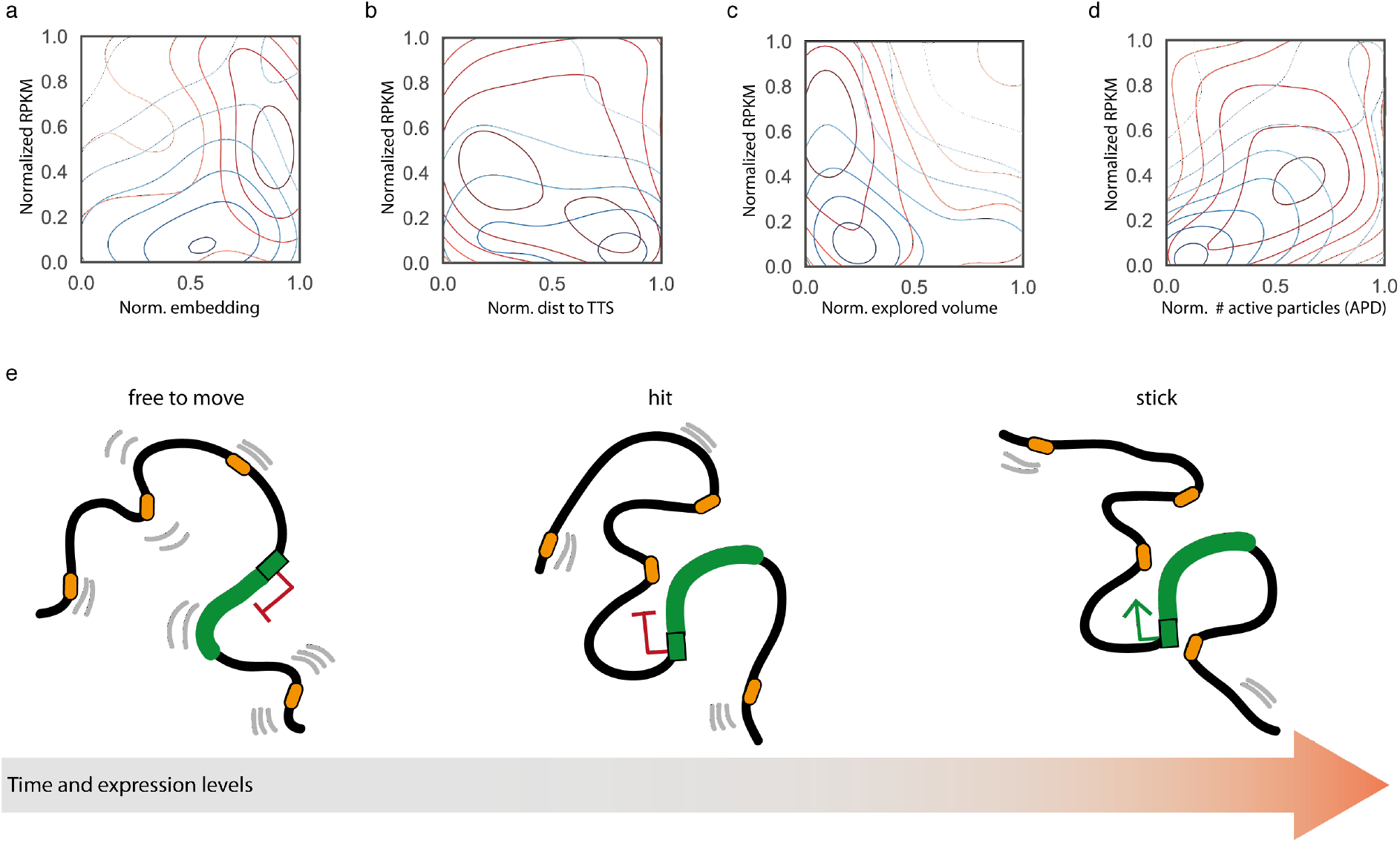
The hit-and-stick model of dynamic gene activation. Density distributions of normalized RPKM expression levels for two sets of simulated loci (that is, continuously active loci in red and loci that switch activity state during simulation in blue) with respect to: **a.** normalized TSS embedding, **b.** normalized distance to TSS, **c.** normalized explored volume, and **d.** normalized number of active APD particles. **e.** Cartoon of the Hit-and-Stick model for gene activation suggested by the TADdyn simulations over 11 *loci*.

In summary, our results support what we call here a *hit-and-stick* model for gene activation. In this model, gene expression is triggered by the formation of a structural domain hosting enhancer sequences (*ADP* particles), with the TSS being *caged* inside the newly formed domain. The proximity to the enhancers and the newly established interactions (*hit*), stabilizes (*stick*) the spatial position of the *locus* TSS against the random thermal motion (**Fig. 5e**).

## DISCUSSION

Here we have introduced a new computational tool, TADdyn, aimed at studying time-dependent dynamics of chromatin domains during a cell fate conversion process that models smooth 3D transitions of chromosome domains. To our knowledge this is the first completely 3C-based data-driven method to simulate structural changes of chromatin over time. TADdyn takes as input a discrete time-series of Hi-C interaction maps and converts matrix bins into spherical particles. These are dynamically and spatially restrained using Hi-C interactions^6^ and a physics-based chromatin model^21^, effectively integrating restraint-based modeling and polymer physics simulations. For our study, we modeled the dynamic rearrangement of 11 loci during the reprogramming of murine B cells to pluripotent stem sells^34^, integrating *in situ* Hi-C data obtained for 7 consecutive stages. Molecular dynamics simulations of data at 5kb resolution predict viable structural and dynamical transitions between the different cell stages in 3D space. TADdyn is therefore suited to model processes characterized by continuous chromatin structural transitions.

In two alternative simulations of *Sox2*, we found that missing steps only marginally affect the results of our simulations suggesting that the simulated reprogramming dynamics are dominated by smooth and gradual spatial chromatin rearrangements. In fact, TADdyn has turned out to be useful in determining whether a smooth spatial reorganization of a chromatin region between two consecutive time steps is physically feasible or is hindered by “energetic barriers” between the structures to be simulated.

TADdyn simulations have provided new insights into mechanisms underlying chromatin domain formation and gene expression regulation, extending our current understanding of these processes. First, we found that upon activation, the promoter region establishes interactions with *active* chromatin particles that contain overlapping signals of chromatin accessibility (ATAC-seq) and regulatory activity (H3K4me2) (**Fig. 4**). These promoter-*active* interactions build a local structural domain that embeds the TSS and makes it significantly less accessible to external particles and more spatially constrained against random thermal motion (**Figs. 3** and **4**). Interestingly, these findings are consistent with previous super-resolution imaging studies suggesting that the mobility of the promoter is constrained upon activation^36,37^.

Notably, these *active* particles with ATAC-seq and H3K4me2 signal often include annotated enhancers as well as newly predicted putative enhancers. Such predicted enhancers can act at long-range, as exemplified by the distal *active* particles interacting with the *Sox2* promoter. Upon transcription activation, the dynamic formation of local topological domains (*cages*) inferred here from high-resolution time-resolved Hi-C data, is reminiscent of structural or topological motifs (such as rosettes, or cliques) suggested in previous studies using polymer-physics-based models^42^ as well as transcription factories or phase-separated condensates^39,40,43,44^. We speculate that these *cages* might trigger and maintain the activation of the promoter by maximizing its local association with a series of proximal enhancers. At a larger scale, such single-gene cages might eventually coalescence into larger domains to form large multi-gene condensates.

Based on our findings that the active promoter becomes *caged* inside a structural domain together with open and active regions we propose a *hit-and-stick* model for promoter-enhancer transcription regulation (**Fig. 5e**). Our model predicts that this molecular *cage* ensures the establishment (*hit phase*) and the maintenance of a spatial neighborhood where promoter-enhancers interactions are favored for the entire active phase (*stick phase*). This second phase is accompanied by low accessibility to external particles and stabilization of the TSS against thermal motion. The model also proposes that promoter-enhancer communication^45^ occurs via direct interactions between the distant enhancer and its cognate target gene as previously observed in many studies^33,37,43,44,46–54^. In summary, our hit-and-stick model links the promoter-enhancer direct contact to the formation of local structural domains or “cages”.

Our results seem incompatible with alternative mechanisms of gene regulation that do not necessarily depend on promoter-enhancer interactions^55,56^ nor its extension in time^57^. In such an alternative scenario, promoter-enhancer contacts are short-lived but continue to maintain a transcriptionally-competent chromatin environment at promoters to ensure their activity even if long-range interactions are lost. However, as the time difference between consecutive reprogramming samples used to obtain Hi-C interaction maps was in the order of tens of hours we cannot rule out that at smaller scales cell sub-populations diverge from the general mechanism here proposed. Therefore, this limitation could have precluded detection of other possible dynamic pathways of genome conformation, satisfying only a subset of the input interaction matrices. To address this, TADdyn will have to be implemented in the future using data obtained from finer time-resolved Hi-C or imaging-based approaches^58,59^.

## METHODS

### Collection of experimental data

Structural data were obtained from 3C-based chromatin interaction experiments previously generated by us^34^. Specifically, *in-situ* Hi-C datasets were downloaded from the GEO database (accession number GSE96611) for the cells stages from B to PSC cells of the reprogramming process in mouse. The dataset included seven Hi-C experiments: B cells (B), Bα cells (after 18h, Bα), day2 (after 48h, D2), day4 (after 48h, D4), day6 (after 48h, D6), day8 (after 48h, D8), and Pluripotent Stem Cells (after about 48h, PSC). The reads were mapped and filtered as previously described^34^. Using these filtered fragments, the genome-wide raw interaction maps were binned at 5 Kilobase (kb) resolution and normalized using “Vanilla” algorithm^60,61^ as implemented in TADbit^62^. Genomic regions were selected around 11 loci of interest (**Table 1**) (*Sox2*, *Nanog*, *Mmp3*, *Mmp12*, *Ebf1*, *C/EBPα*, *Rad23a*, *Rad23b*, *Lmo7*, *Nos1ap* and *Tet2*) each characterized by a specific gene expression profile during the cell reprogramming process (**Supplementary Figs. 1-11a**). The majority of the genes (*Sox2*, *C/EBPα*, *Rad23a*, *Rad23b*, *Tet2*) span less than 100kb and where simulated in the center of Megabases (Mb) chromatin regions. For the *Nanog locus*, the modelled region contained only 1Mb around the *locus* after filtering low coverage bins (that is, those with more than 75% of cells with zero counts), and for the *Mmp12* (chr9:7,344,381-7,369,499) and *Mmp3* (chr9:7,445,822-7,455,975) *loci* that are contained in the region a single region of 1.5Mb was considered (chr9:6,650,000-8,150,000). The remaining loci (*Ebf1*, *Lmo7*, *and Nos1ap*) were longer than 100kb and were simulated in 3.0Mb regions centered on the promoter (**Fig. 1a** and **Supplementary Figs. 1-11b**). The possibility to study these regions using restraint-based modelling was tested *a priori* computing the matrix modelling potential (MMP)^35^ on the 70 normalized sub-matrices (that is, the 10 genomic regions and the 7 cell stages). In brief, all Hi-C interaction matrices had MMP scores higher than 0.78, and contained very few low-coverage bins (<0.05% of the bins), indicating that the restrained-based approach at 5kb should provide accurate 3D models for all the regions^35^ (**Supplementary Figs. 1-11b**).

### Representation of the chromatin region using bead-spring polymer model

Chromatin was represented as a bead-spring polymer describing the effective physical properties of the underlying chromatin^18,21^. Specifically, each Hi-C matrix bin of 5kb was represented as a spherical particle of diameter 50 nanometers (nm) using a compaction ratio of 0.01bp/nm^63,64^. Additionally, two non-harmonic potentials were introduced taking into account the *excluded volume* interaction (purely repulsive Lennard-Jones) and the *chain connectivity* (Finitely Extensible Nonlinear Elastic, FENE). Both potentials were used as previously described^18^. In the present application, the *bending rigidity* potential, although it is available in TADdyn, is not used for consistency with the initial models generated using the TADbit software at the B stage (see below). The center of mass of the chromatin chain was tethered to the origin *O*=(0.0,0.0,0.0 x, y, z coordinates, respectively) of the system using a Harmonic (K_t_=50., d_eq_=0.0), and was simulated inside a cubic box of size 50μm (much larger of the size of the models), which was also centered at *O*.

### Encoding of the experimental data into TADdyn restraints

TADdyn accepts one of three possible initial conformations for the chromatin: (i) a polymer chain prepared as a random walk, (ii) an arrangement made of stacked rosettes^18^, or (iii) a previously generated data-driven model. For the 100 time-series simulations performed here, the initial conformations were the 100 optimal models were built using TADbit (https://3DGenomes.org/tadbit) as previously described^62^ for the B cell time point (the first stage of the Hi-C time series). Specifically, we explored all possible combinations of the parameters (*lowfreq*, *upfreq*, *maxdist*, *dcutoff*)^65^ in the intervals *lowfreq*=(−3.0,−2.0,−1.0,0.0), *upfreq*=(0.0,1.0,2.0,3.0), maxdist=(150,200,250,300,350,400)nm, and *dcutoff*=(150,175,200,225,250)nm. To select the optimal set of parameters, we computed the correlation of the input Hi-C interaction map in the B cell stage and the models contact map at the different distance cutoffs using only the 100 best (that is, with the lowest objective function) models per each triplet of the 500 generated models. Next, per each triplet of parameters, the median Spearman correlation values were computed over the 11 studied loci. The largest median correlation coefficient of 0.78 was obtained for *lowfreq=1.0*, *upfreq=1.0*, *maxdist*=300nm, and *dcutoff* =225nm.

Next, the 100 TADbit generated models in B cells were energy minimized using a short run of the Polak-Ribiere version of the conjugate gradient algorithm^66^ to favor smooth adaptations of the implementations of the *excluded volume* and *chain connectivity* interaction in TADdyn.

The optimal TADbit parameters optimized for the B stage (that is, *lowfreq* of −1.0, *upfreq* of 1.0, and *maxdist* of 300nm) were then used to define the set of distance harmonic restraints of the other time points of the series (**Fig. 1b**). To adapt the harmonic restraint of a given pair of particles *(i,j)* between consecutive time points *t*_*n*_ and *t*_*n+1*_, one of the following 3 possible scenarios was applied (**Fig. 1c**):

1. If the pair *(i,j)* was restrained by the same type of distance restraint (Harmonic or LowerBoundHarmonic) in both *t*_*n*_ and *t*_*n+1*_, the strength (*k)* and the equilibrium distance (*d*_*eq*_) of the harmonic were both changed gradually from the values they had in *t*_*n*_ to the values they had in *t*_*n+1*_.
2. (a) If the distance restraint applied between *(i,j)* was present at time *t*_*n*_, but vanished at time *t*_*n+1*_, the strength *k* was decreased from the value at *t*_*n*_ to 0.0, and the equilibrium distance *d*_*eq*_ was kept constant and equal to the value in *t*_*n*_. (b) If the distance restraint was present only at time *t*_*n+1*_, the strength *k* was increased from 0.0 at *t*_*n*_ to the value in *t*_*n+1*_, and the equilibrium distance *d*_*eq*_ was kept constant and equal to the value in *t*_*n+1*_.
3. If the pair *(i,j)* was restrained by different type of distance restraint (Harmonic to LowerBoundHarmonic, or vice versa) in *t*_*n*_ and *t*_*n+1*_, two distance restraints were defined for *(i,j)*. The restraint which was active at time *t*_*n*_ was then switched off as in case 2a, and the one active at time *t*_*n+1*_ was switched on as in 2b.

### TADdyn restraint-based dynamics simulations

By applying the previous protocol, the simulation effectively and smoothly modified the underlying restraints during the steered transition from *t*_*n*_ to *t*_*n+1*_. The dynamics of the system was thus described using the stochastic (Langevin) equation^67^, which was integrated using LAMMPS^68^ (http://lammps.sandia.gov) with values of the particle mass (*m=1.0*), the friction (γ=0.5 τ_LJ_^−1^), and the integration time step of *dt*=0.001 *τ*_*LJ*_, where *τ*_*LJ*_ is the internal time unit^69^. The time-dependent Harmonic and LowerBoundHarmonic restraints were implemented using the Colvars plug-in for LAMMPS^70^ originally introduced for advanced sampling techniques, and here modified to implement the TADdyn transitional restraints. This modified version is freely available at https://github.com/david-castillo/colvars. The transition between consecutive time stages was set to last for 10 *τ*_*LJ*_, hence a single run to simulate the complete B to PSC reprogramming process passing through the 7 cell stages lasted for 60 *τ*_*LJ*_. In each of the 100 replicates of the reprogramming run, the model conformations were stored every 0.1 *τ*_*LJ*_, and used for further data analysis. To test the predictive power of TADdyn in case of smooth structural rearrangements two additional simulations were performed for the Sox2 *locus* by removing the restraints of stages D2 (ΔD2) and D6 (ΔD6), and by extending the corresponding transitions (Bα → D4 and D4 → D8 respectively) to keep constant the total duration of the runs.

#### Analysis of TADdyn models

##### Time dependent contact maps and TADdyn models assessment

(i) The contact maps were computed at each time step based on the probability within the 100 simulations that pairs of particles are contacting (that is within a distance cut-off of 200nm) (**Fig. 2a** and **Supplementary Figs. 1-11c**). (ii) The resulting contact maps along the simulations were clustered. The Spearman’s rank correlation coefficients (SCC) was computed with each of the time-series Hi-C interaction map. The SCCs were converted into normalized distances from 0 (max SCC) to 1 (min SCC) and used to cluster the contact maps using the Ward hierarchical approach criterion^71^ as implemented in R (**Fig. 2b** and **Supplementary Figs. 1-11d**). (iii) For each simulation, the Root Mean Square Deviation (RMSD) between the optimally superimposed models (separated by 1*τ*_*LJ*_) was computed. Next, per each model pair, the median dRMSD over the 100 replicates was computed. Finally, the structural clustering analysis was done on the matrix of median dRMSDs by using the Ward hierarchical approach criterion as implemented in R (**Fig. 2c** and **Supplementary Figs. 1-11e**).

##### Time dependent measures of the TADdyn structures

(i) The accessibility (A) of each particle in the ensemble of models was calculated using TADbit with parameters nump=100, radius=50nm, and super-radius=200nm. The accessibility (A) transformed into *embedding* (E=1.0-A) that is a measure of the propensity of a particle to be caged inside an internal cavity. The *embedding* ranges from 0.0 meaning that the particle is located on the interface of the model to 1.0 when the particle is closely surrounded by other particles inside the model (*caged*) (**Fig. 3a** and **Supplementary Figs. 1-11f**). (ii) The explored volume per particle every 5 *τ*_*LJ*_ of trajectory was calculated. Specifically, all trajectories were partitioned in time intervals of 5 *τ*_*LJ*_. In each time interval, the 50 positions (*i.e.* one every 0.1 *τ*_*LJ*_) occupied by the particle *i* were considered, and the convex-hull embedding these 50 positions was calculated. In a given time interval, the average (over the 100 replicate runs) convex-hull volume explored by particle *i* was considered as the typical volume explored by the model particle *i* during the time interval (**Fig. 3b** and **Supplementary Fig.s 1-11g**). (iii) The spatial distance between selected pairs of particles was computed for the ensemble of simulations as the Euclidian distance in nanometers using TADbit (**Fig. 4a** and **Supplementary Figs. 1-11h**).

##### Time dependent insulation score analysis of TADdyn models

To study the partitioning of the models into structural domains (reminiscent of TADs^72–74^), the insulation score (I-score) analysis^38^ was performed on the models contact maps using the command line --is 100000 --ids 50000 --ez --im mean --nt 0.1 --bmoe 3. The called domain borders, whose border strength was deemed significant by the I-score pipeline, were used for further analysis (**Fig. 4b** and **Supplementary Fig.s 1-11i**).

##### Active particles analysis

In each cell stage, we classified some model particles (5kb) into one of 3 possible categories (**Supplementary Table 2**, **Fig. 4c** and **Supplementary Figs. 1-11l and m**): *Active* (A) are particles hosting at least one overlapping 250bp-peak of ATAC-seq and H3K4me2 (ATAC-seq and H3K4me2 data were obtained from our previous work^34^), *Active-Proximal* (AP) are *Active* particles that are close to the TSS particle of the gene either in absolute terms (spatial distance <200nm) or in relative terms (spatial distance < half of average distance at the genomic separation) for at least half of the duration of the stage. *Active-Proximal-Domain* (ADP) are AP particles that are inside the local domain containing the TSS (see insulation score analysis above).

The statistical comparison between the distributions of the different quantities analyzed (spatial distance, embedding, and explored volume) was performed by bootstrapping (re-sampling with replacement). The re-sampling procedure is repeated 1,000 times for a fixed number of elements, specifically 200 for the comparisons in Fig. 4c and 1000 for the ones in Fig. 5a-c). For the 1,000 comparisons of the re-sampled distributions, we tested for significant differences a Wilcoxon test using R. The higher p-value was assigned to the corresponding pair of distributions under comparison. The comparisons that resulted in maximum p-values<0.01 were deemed to be significantly different.

## DATA AVAILABILITY

The TADdyn approach is currently available as part of the TADbit Github repository (https://github.com/david-castillo/TADbit/tree/poly). The modified version of the COLVAR plug-in for LAMMPS is available at the Github repository (https://github.com/david-castillo/colvars)

## ACKNOWLEDGEMENTS

We thank all the current and past members of the Marti-Renom lab for their continuous discussions and support to the development of TADdyn.

