## Supplementary figures and images for "Dynamic simulations of transcriptional control during cell reprogramming reveal spatial chromatin caging"

### Supplemental figure 1

#### CEBPa chr7:35119293-35121931. Forward

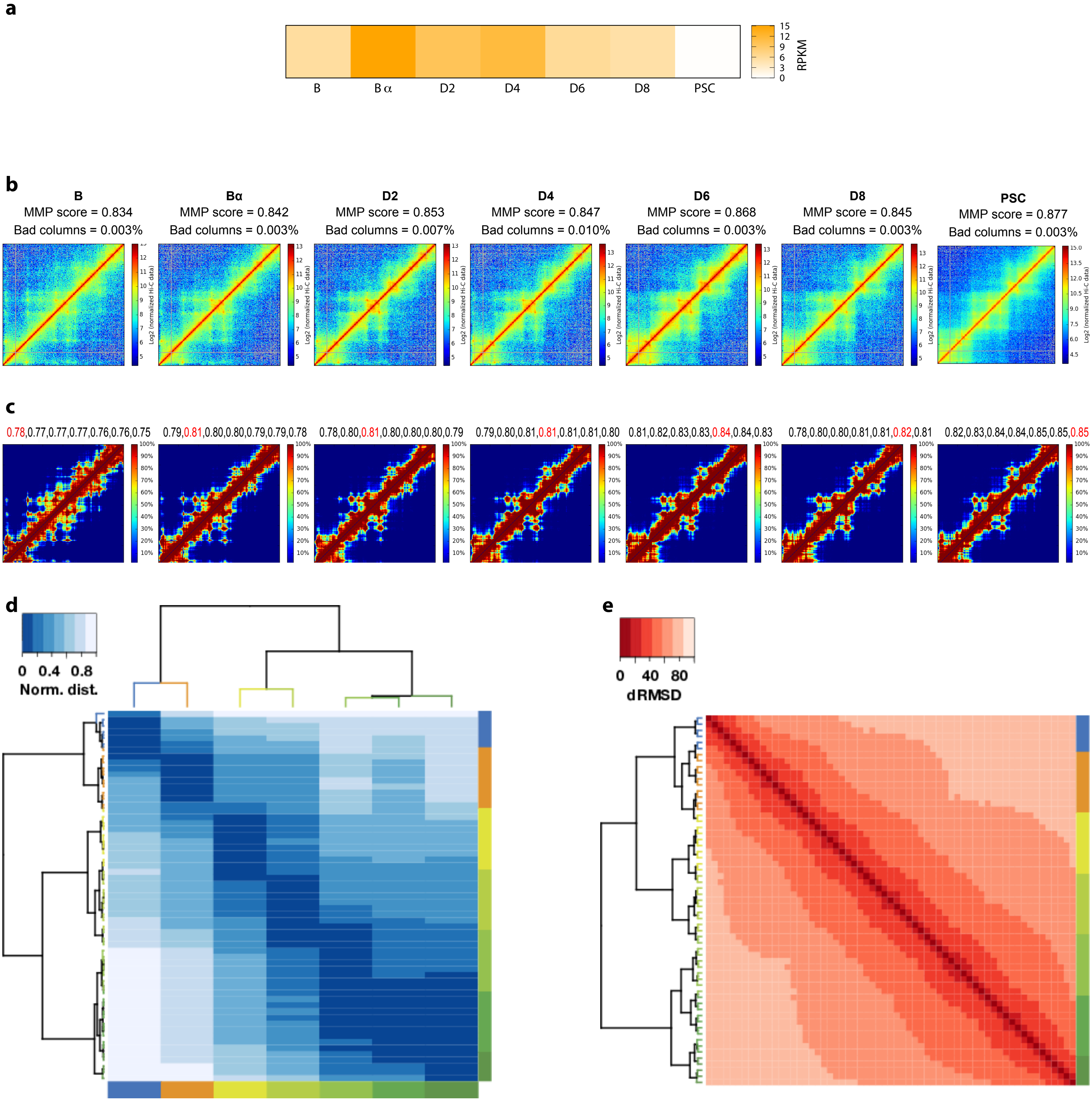

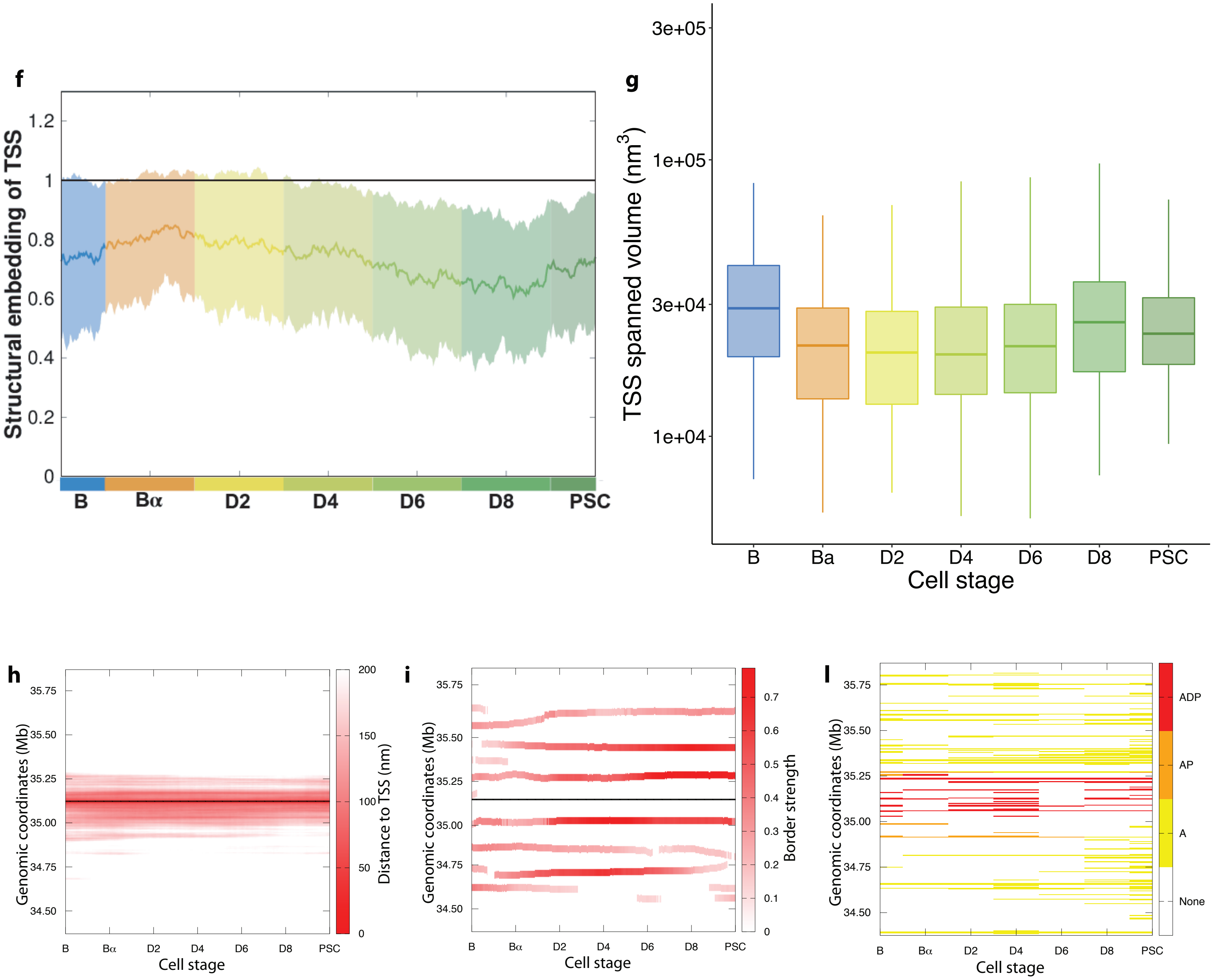


m

| TSS | B | Bα | D2 | D4 | D6 | D8 | PSC |
|---|---|---|---|---|---|---|---|
| **A** | 45 | 33 | 41 | 58 | 36 | 54 | 85 |
| **AP** | 18 | 12 | 14 | 19 | 6 | 9 | 12 |
| **APD** | 10 | 7 | 11 | 16 | 5 | 9 | 11 |

### Supplemental figure 2

#### Ebf1 chr11:44618100-45008096. Forward

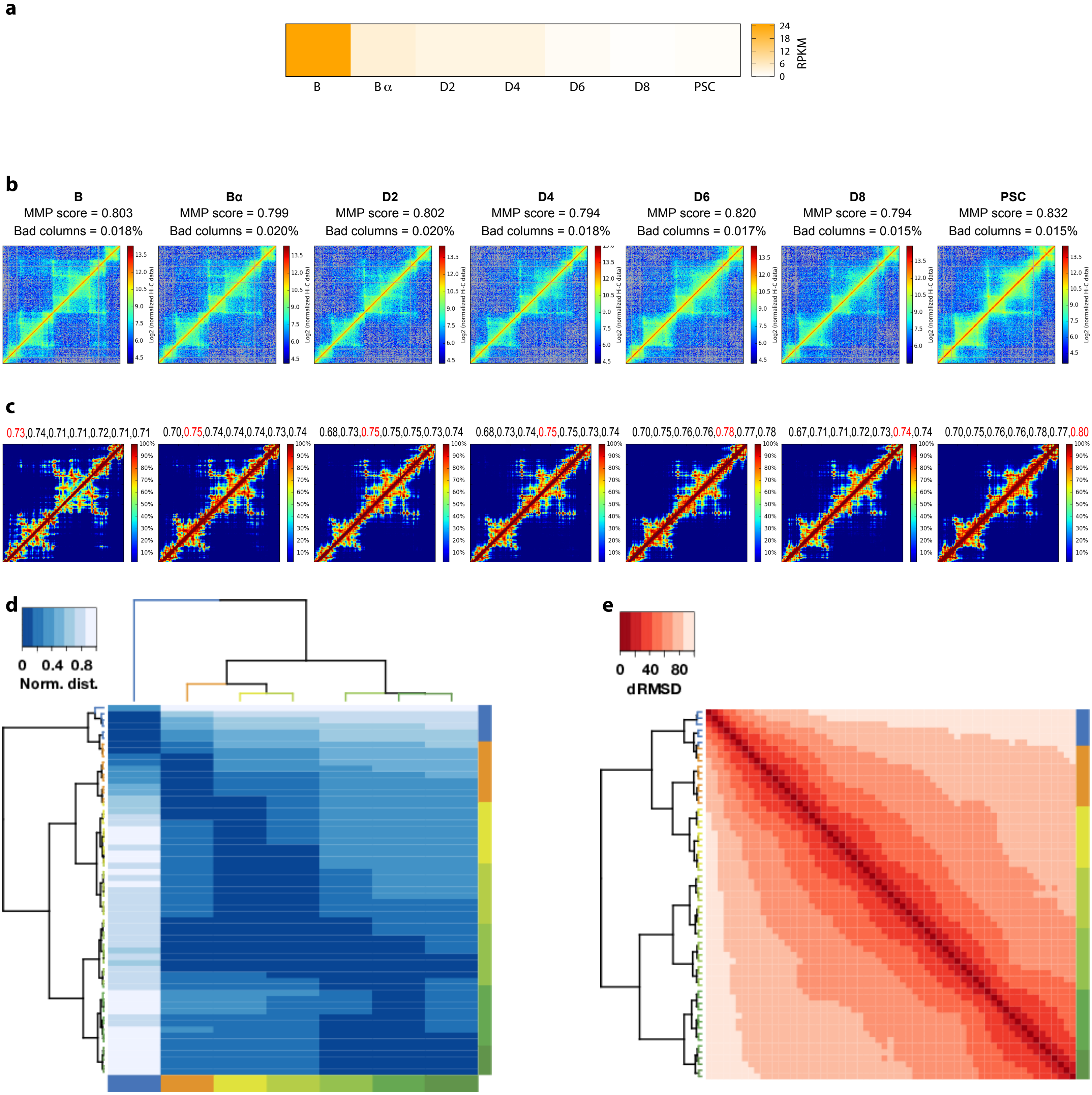

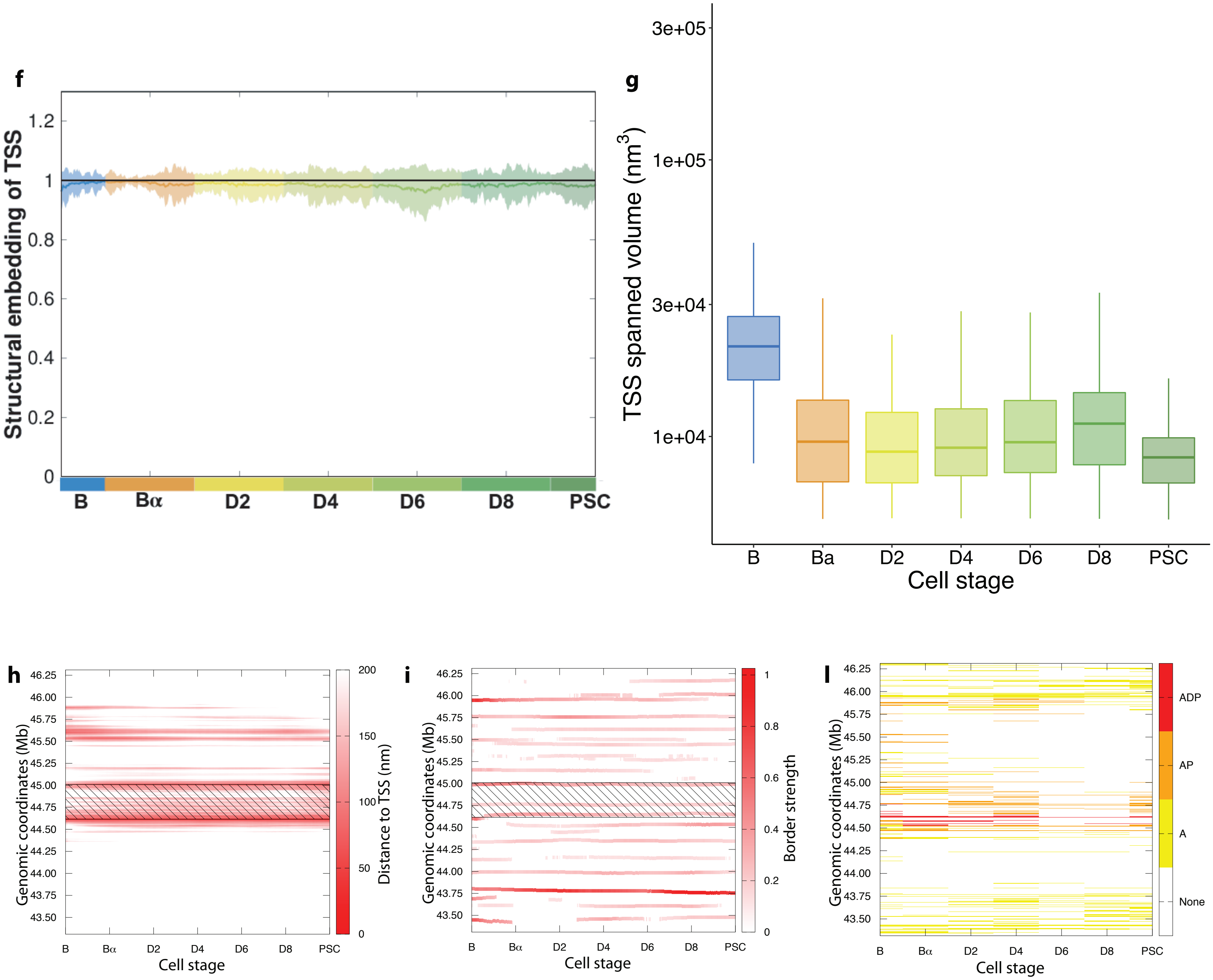


**m**

| TSS | B | Bα | D2 | D4 | D6 | D8 | PSC |
|---|---|---|---|---|---|---|---|
| **A** | 106 | 85 | 78 | 101 | 53 | 64 | 88 |
| **AP** | 41 | 35 | 31 | 40 | 13 | 10 | 27 |
| **APD** | 7 | 8 | 5 | 6 | 1 | 2 | 3 |

### Supplemental figure 3

#### Lmo7 chr14:101729957-101934710. Forward

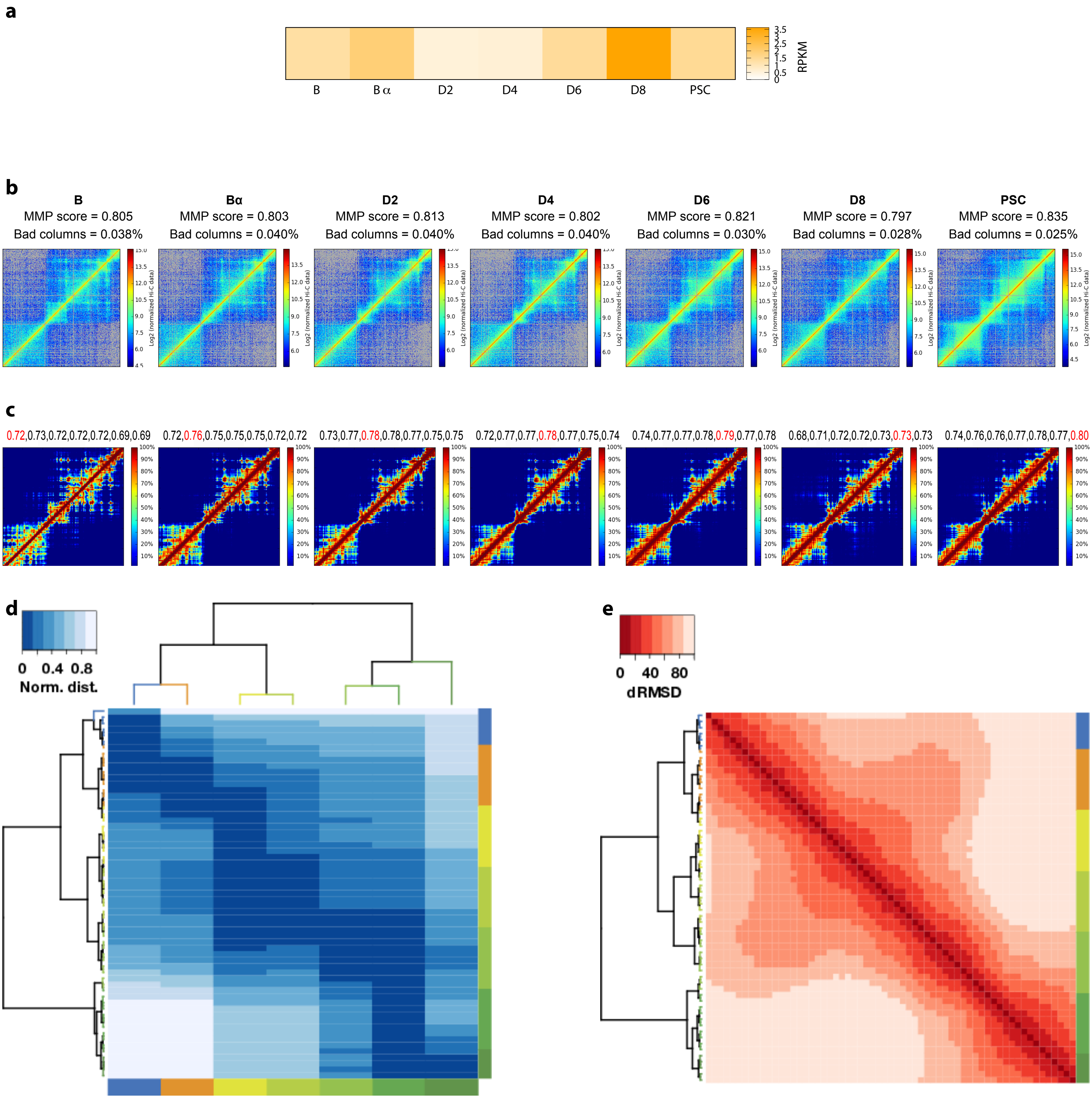

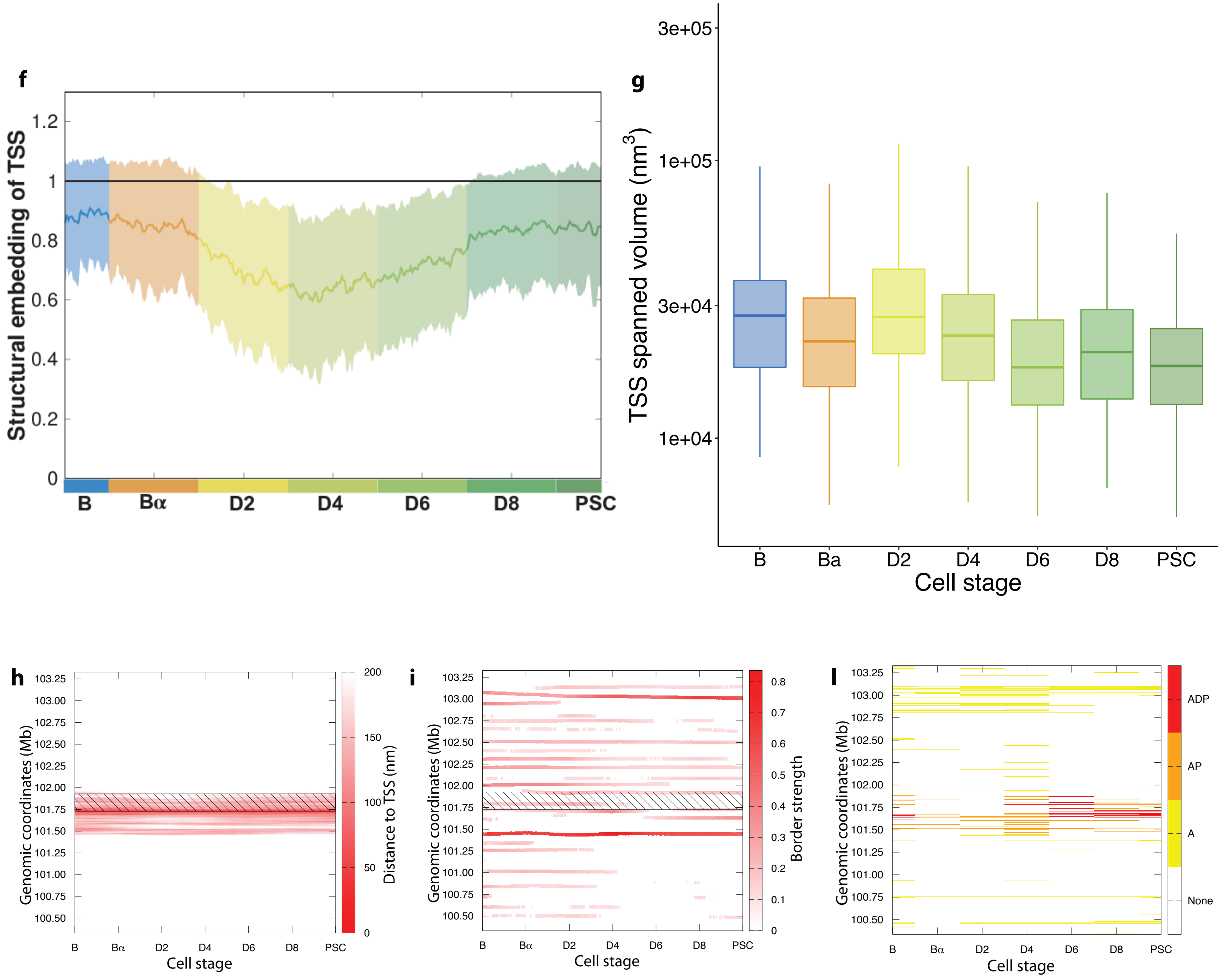


m

| TSS | B | Bα | D2 | D4 | D6 | D8 | PSC |
|---|---|---|---|---|---|---|---|
| A | 58 | 36 | 47 | 70 | 41 | 42 | 44 |
| AP | 19 | 10 | 14 | 28 | 20 | 23 | 21 |
| APD | 7 | 1 | 1 | 2 | 15 | 9 | 10 |

### Supplemental figure 4

#### Mmp3 chr9:7445822-7455975. Forward

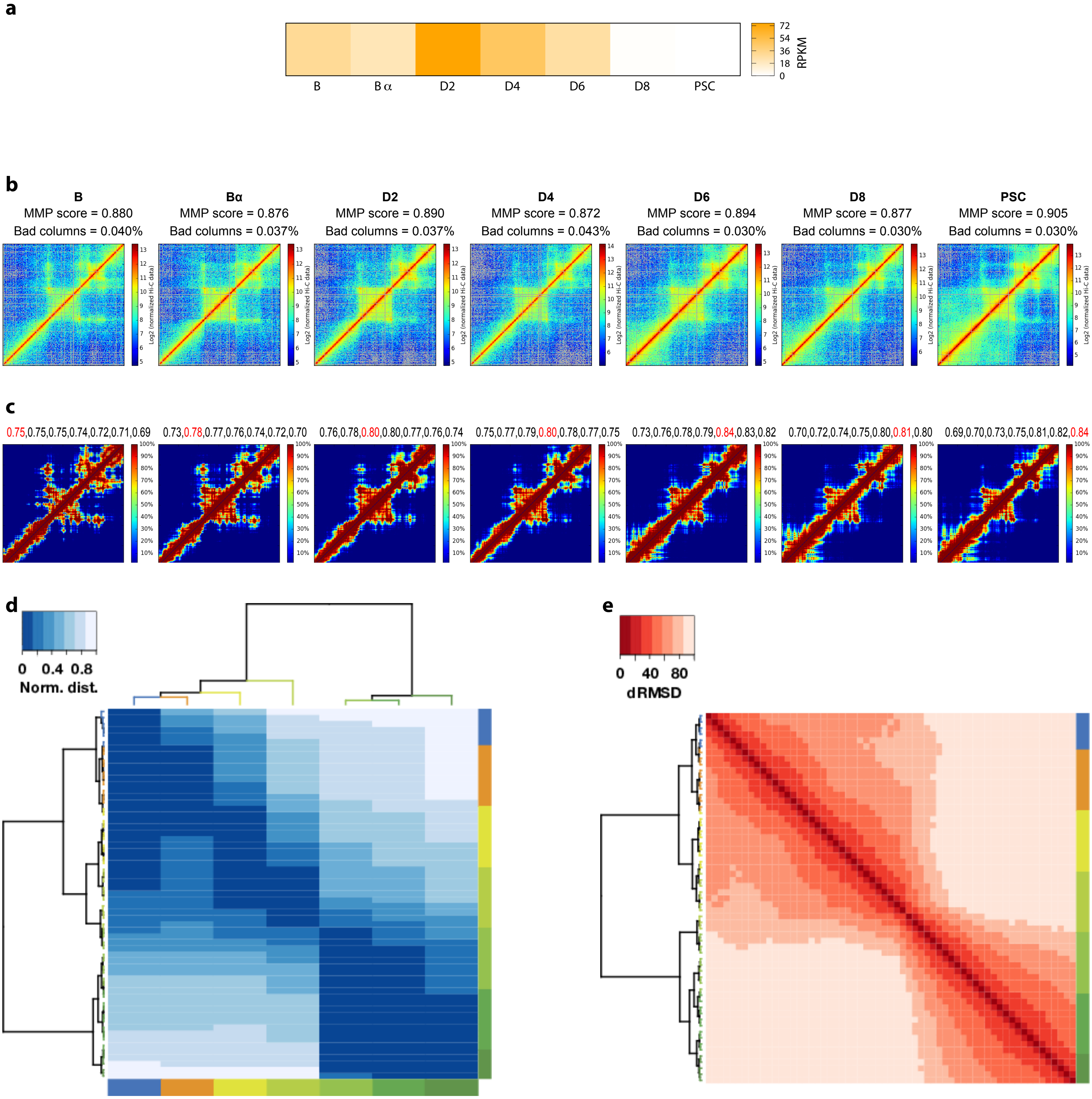

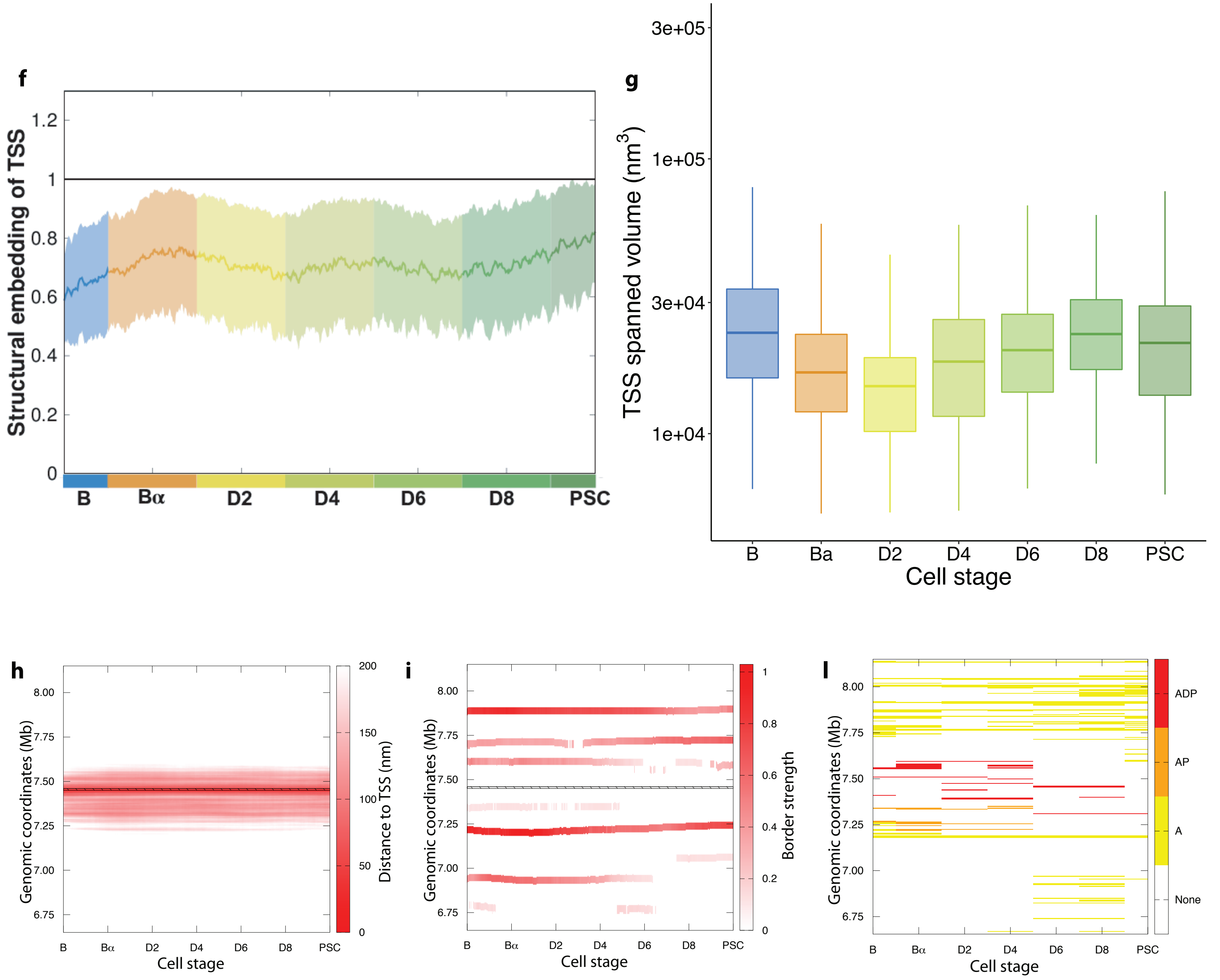


**m**

| TSS | B | Bα | D2 | D4 | D6 | D8 | PSC |
|---|---|---|---|---|---|---|---|
| **A** | 43 | 39 | 25 | 35 | 34 | 48 | 43 |
| **AP** | 7 | 16 | 9 | 12 | 3 | 4 | 1 |
| **APD** | 4 | 8 | 6 | 8 | 3 | 4 | 1 |

### Supplemental figure 5

#### Mmp12 chr9:7344381-7369499. Forward

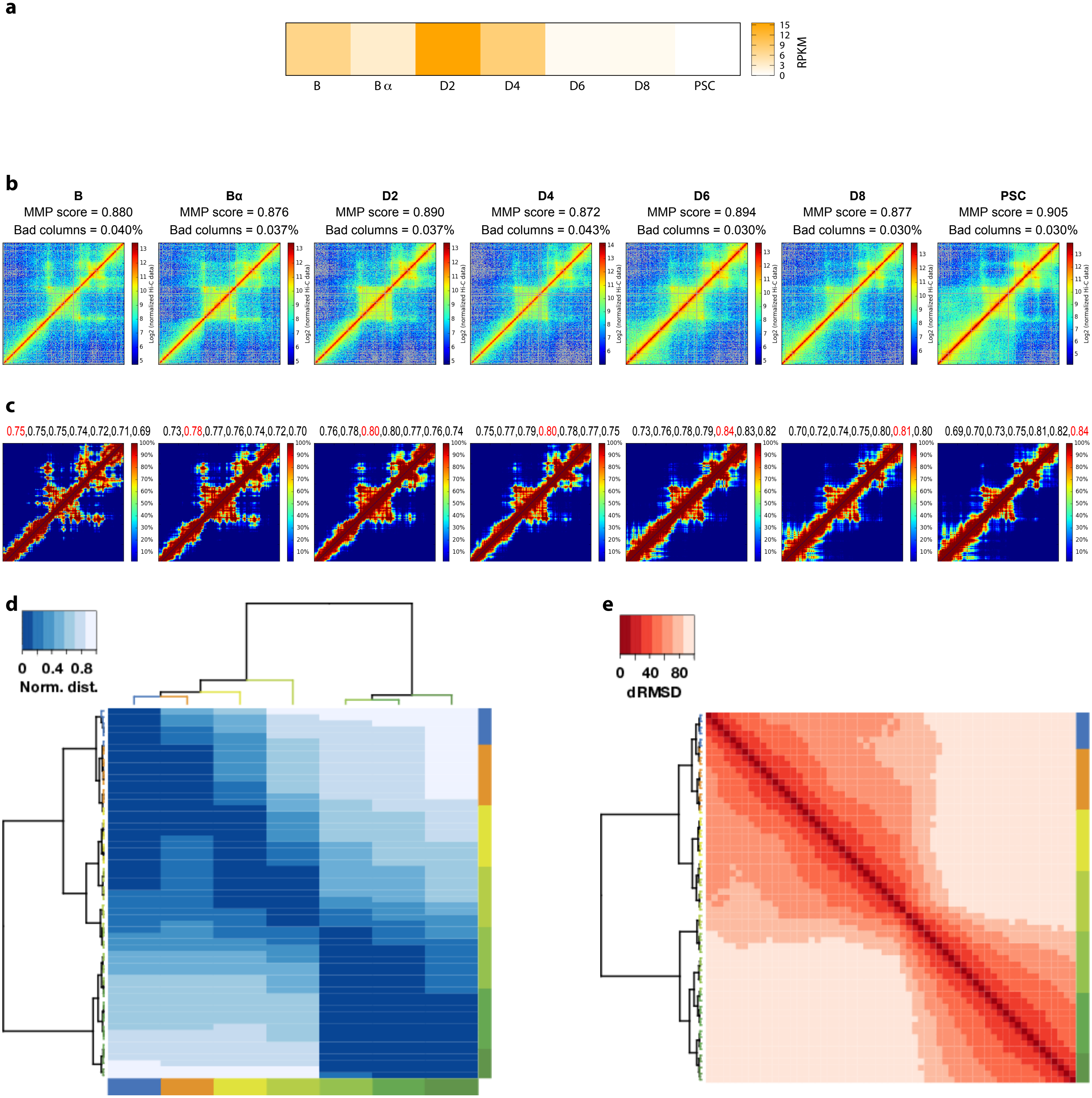

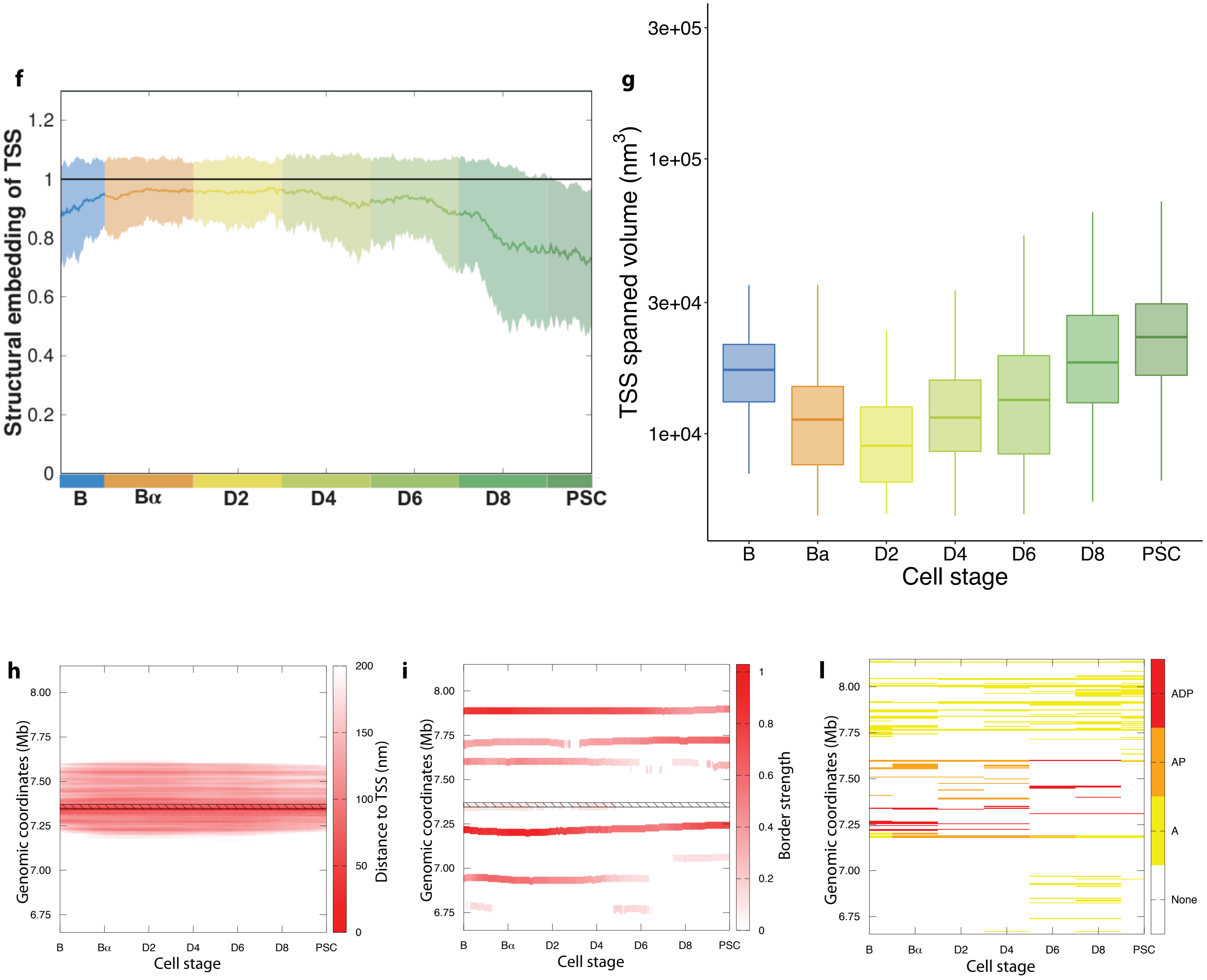


**m**

| TSS | B | Bα | D2 | D4 | D6 | D8 | PSC |
|---|---|---|---|---|---|---|---|
| A | 45 | 40 | 26 | 36 | 36 | 49 | 43 |
| AP | 14 | 22 | 13 | 16 | 7 | 5 | 2 |
| APD | 8 | 8 | 3 | 4 | 5 | 5 | 1 |

### Supplemental figure 6

#### Nanog chr6:122707565-122714633. Forward

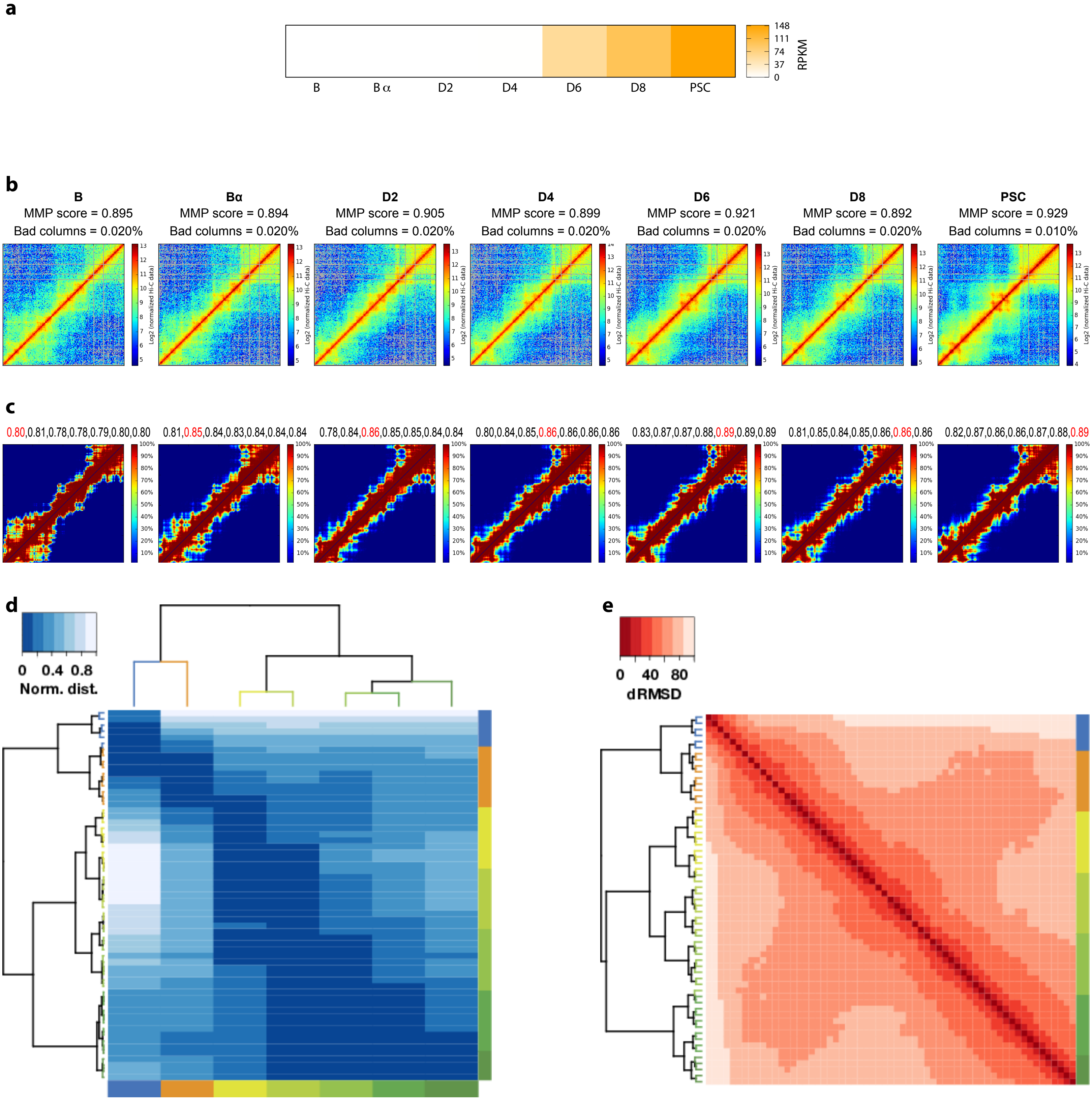

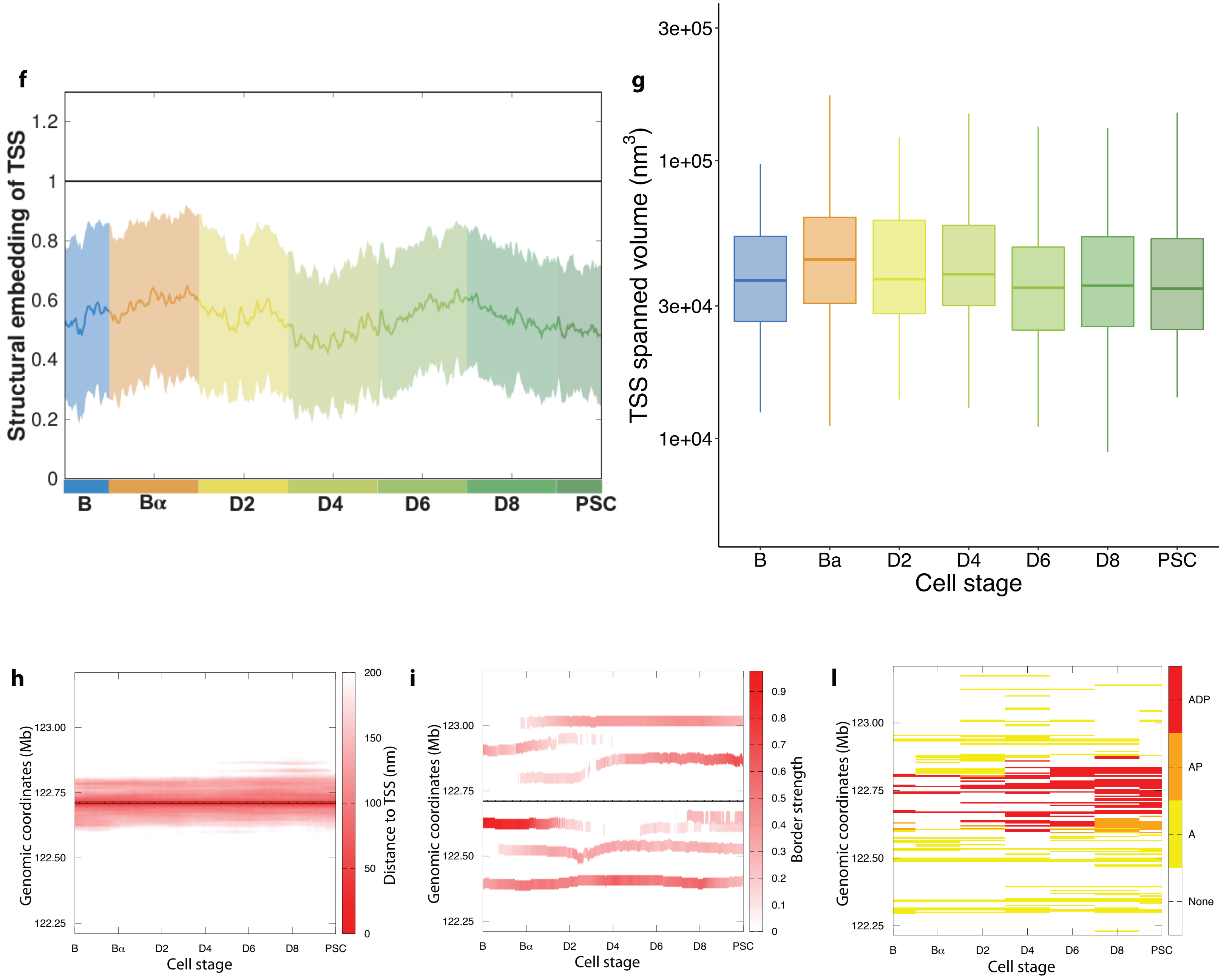


**m**

| TSS | B | Bα | D2 | D4 | D6 | D8 | PSC |
|---|---|---|---|---|---|---|---|
| **A** | 30 | 32 | 47 | 54 | 51 | 68 | 62 |
| **AP** | 11 | 5 | 14 | 23 | 25 | 36 | 36 |
| **APD** | 8 | 5 | 11 | 23 | 23 | 27 | 27 |

### Supplemental figure 7

#### Nos1ap chr1:170302668-170589861. Reverse

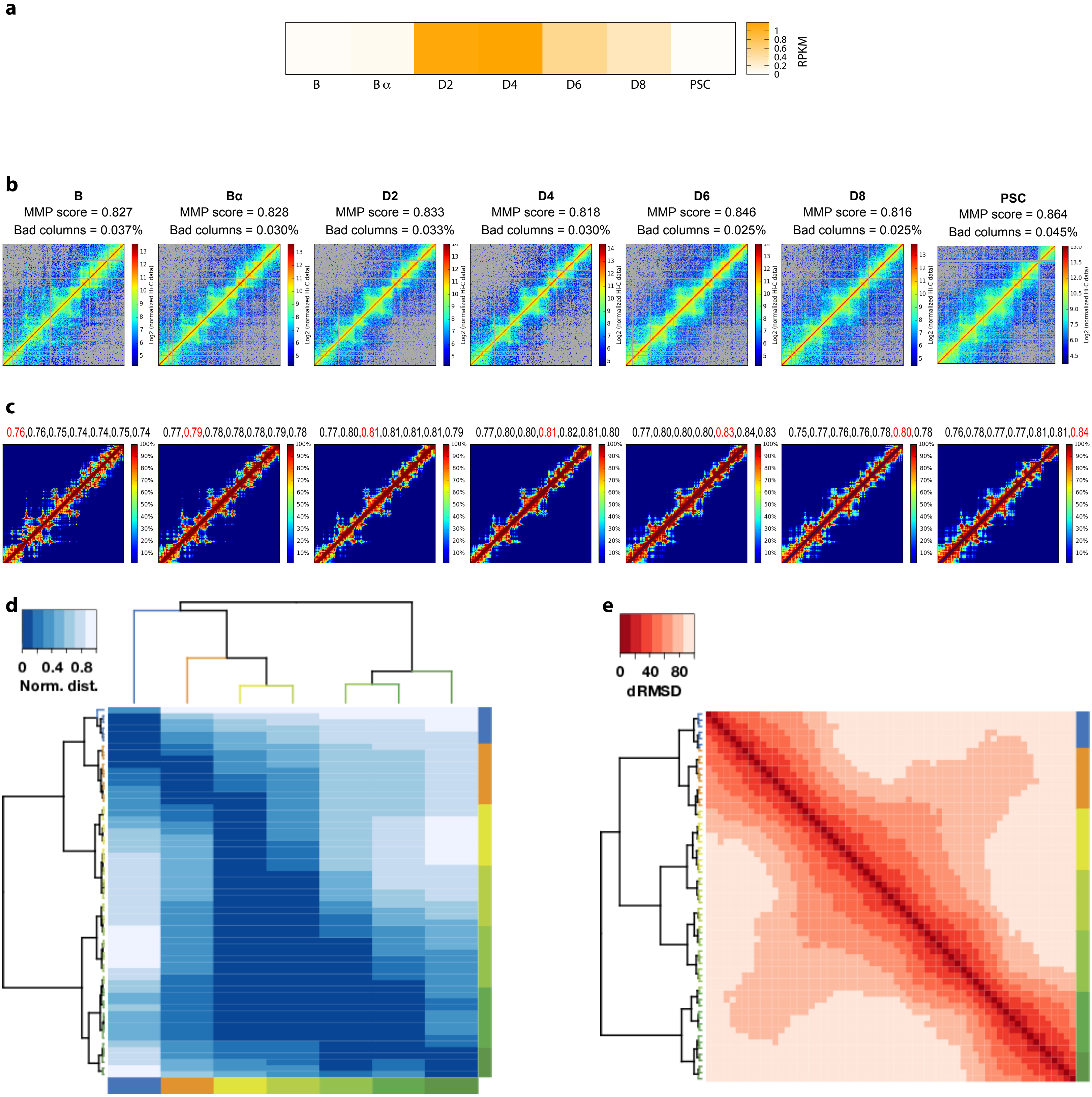

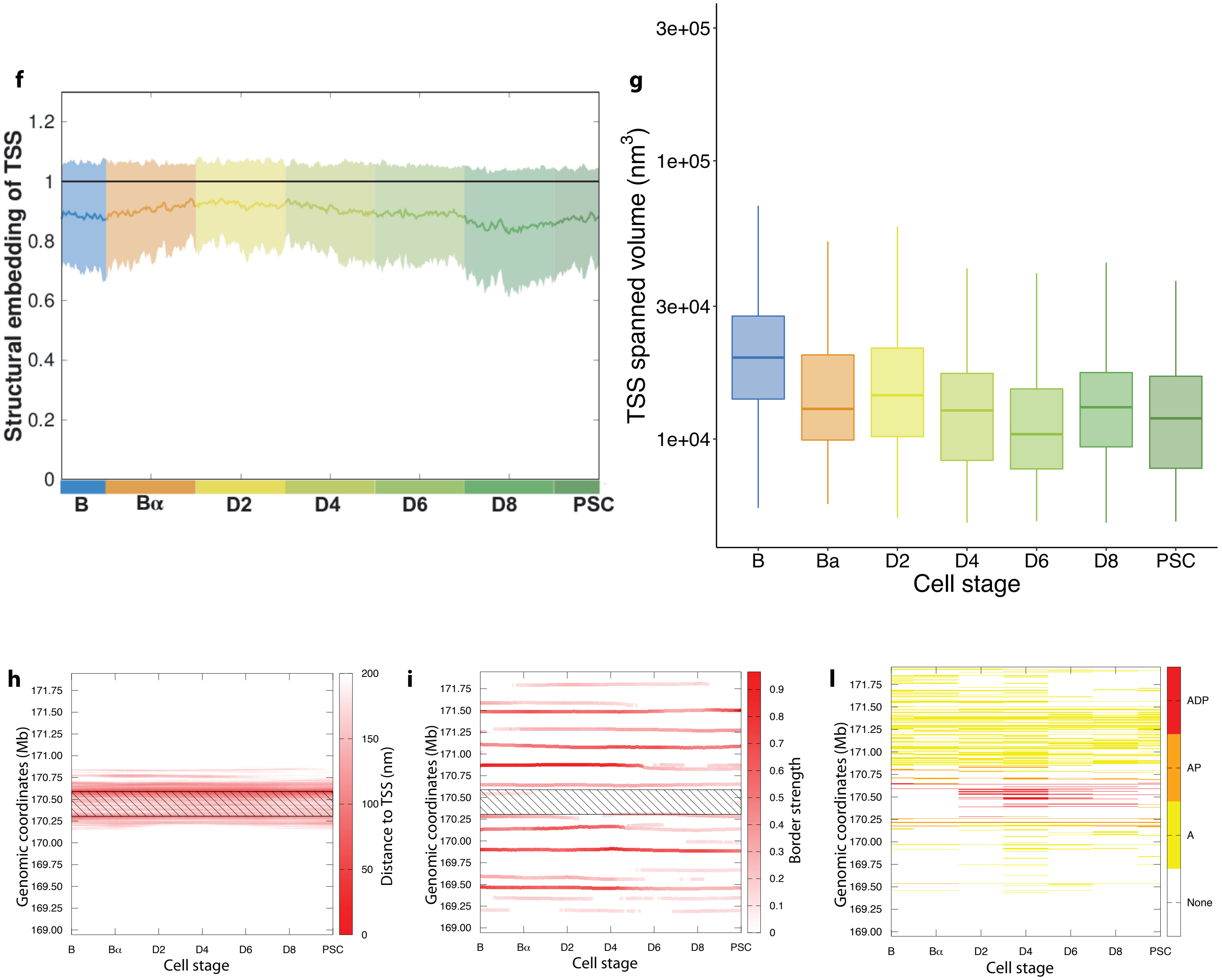


**m**

| TSS | B | Bα | D2 | D4 | D6 | D8 | PSC |
|---|---|---|---|---|---|---|---|
| **A** | 113 | 100 | 108 | 144 | 100 | 93 | 96 |
| **AP** | 16 | 13 | 21 | 32 | 19 | 18 | 16 |
| **APD** | 3 | 2 | 14 | 22 | 13 | 9 | 4 |

### Supplemental figure 8

#### Rad23a chr8:84834652-84840665. Reverse

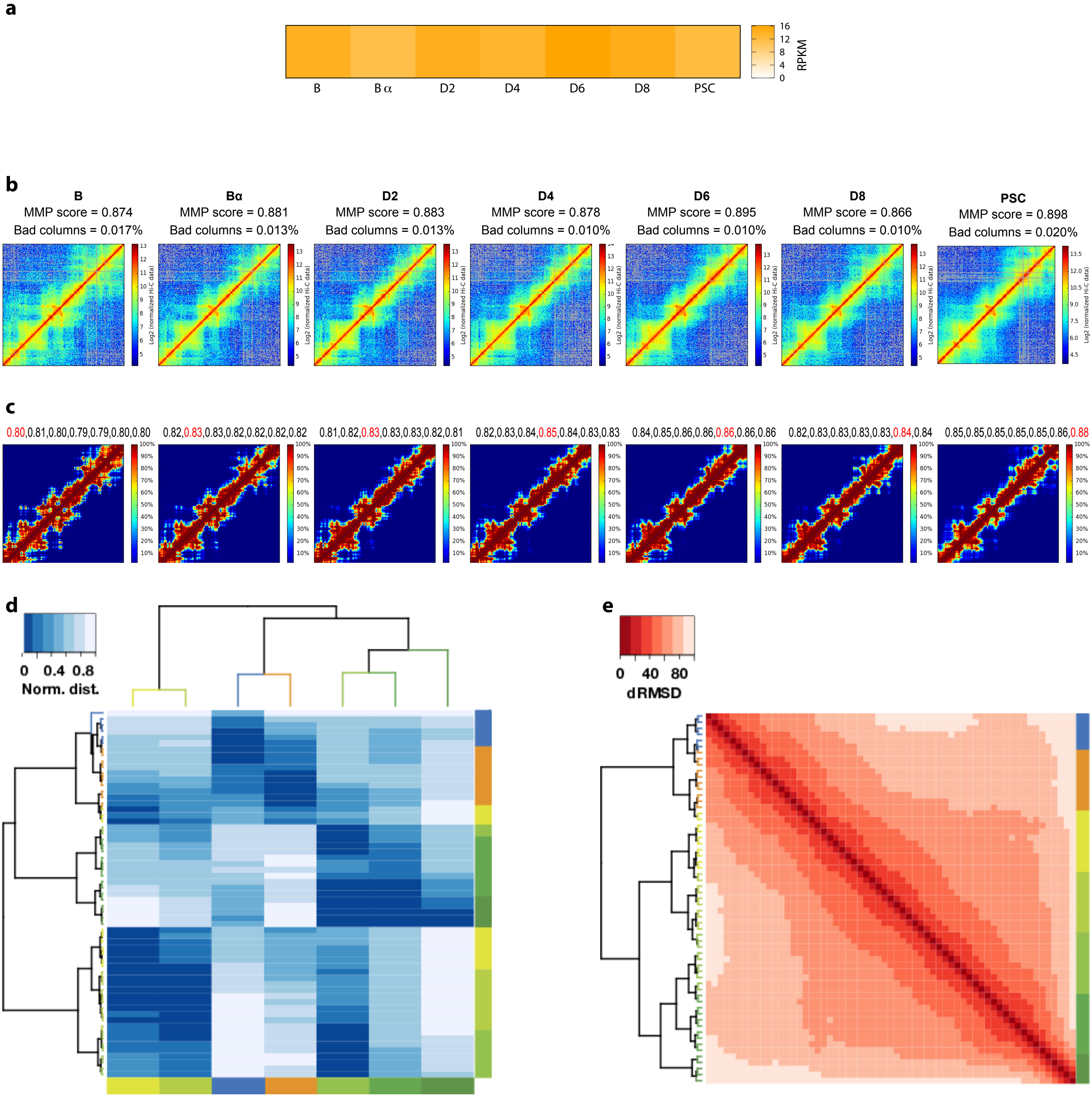

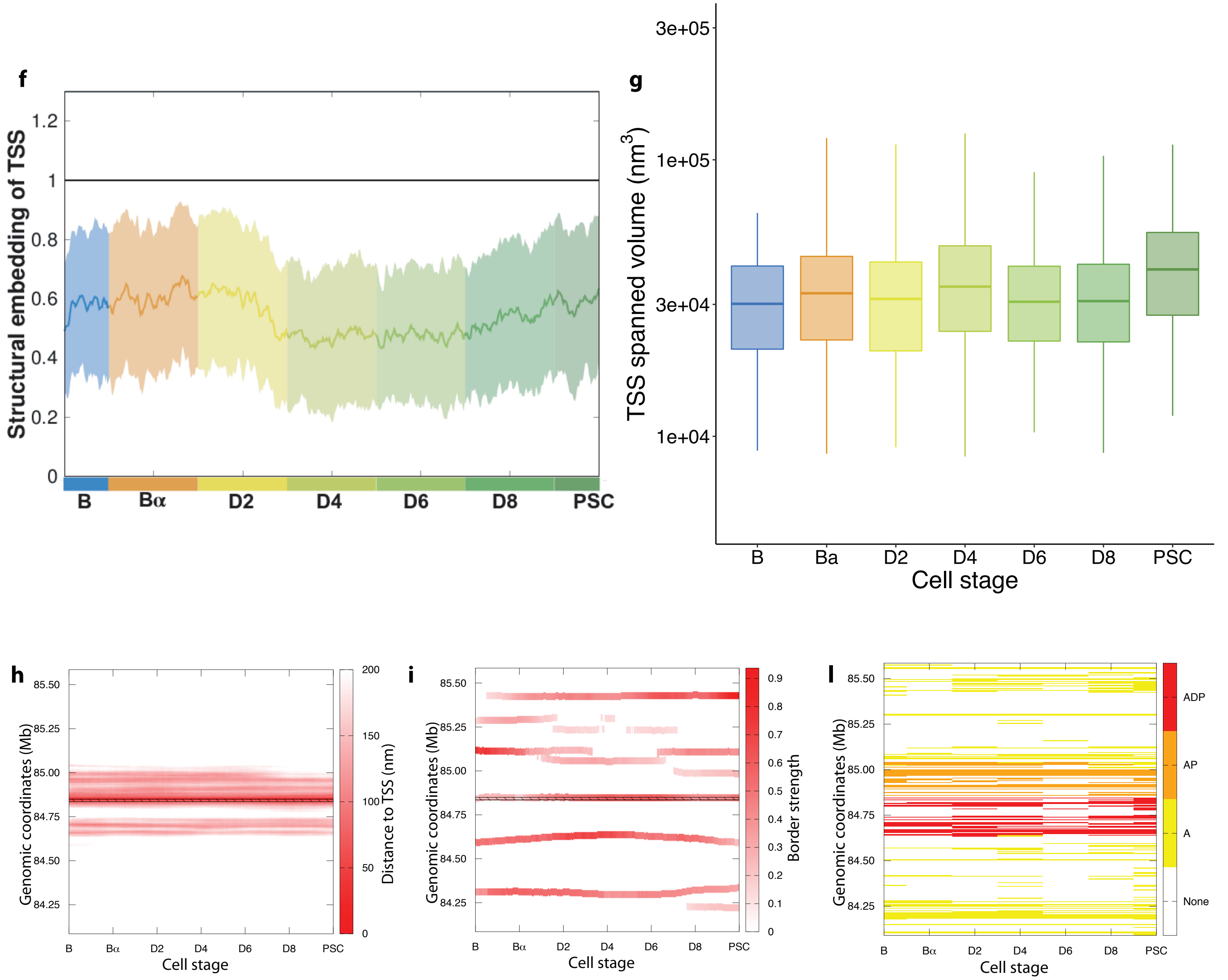


**m**

| TSS | B | Bα | D2 | D4 | D6 | D8 | PSC |
|---|---|---|---|---|---|---|---|
| **A** | 82 | 76 | 87 | 95 | 67 | 92 | 119 |
| **AP** | 41 | 40 | 46 | 43 | 31 | 38 | 48 |
| **APD** | 18 | 19 | 25 | 21 | 14 | 20 | 28 |

### Supplemental figure 9

#### Rad23b chr4:55350042-55392237. Forward

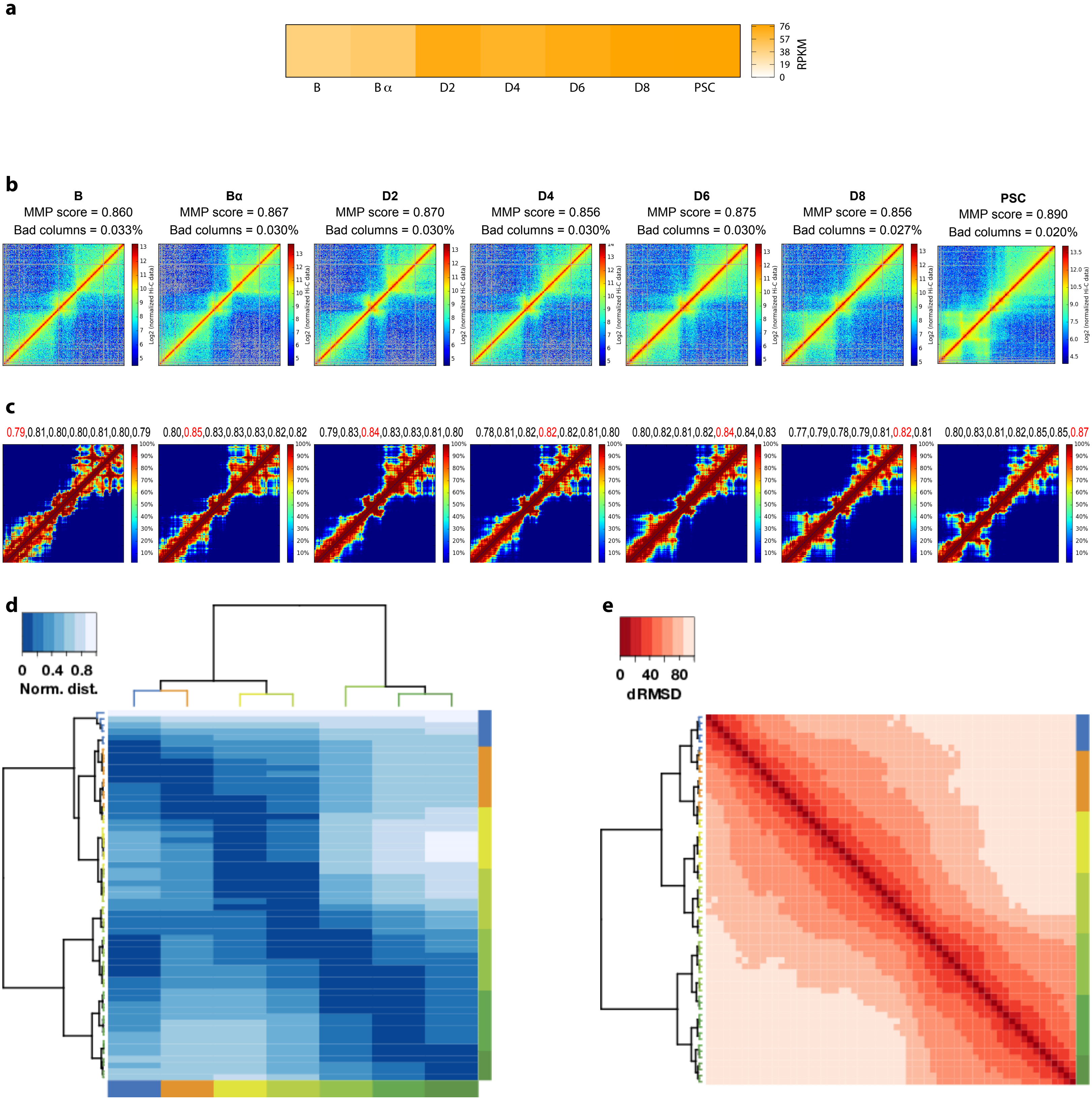

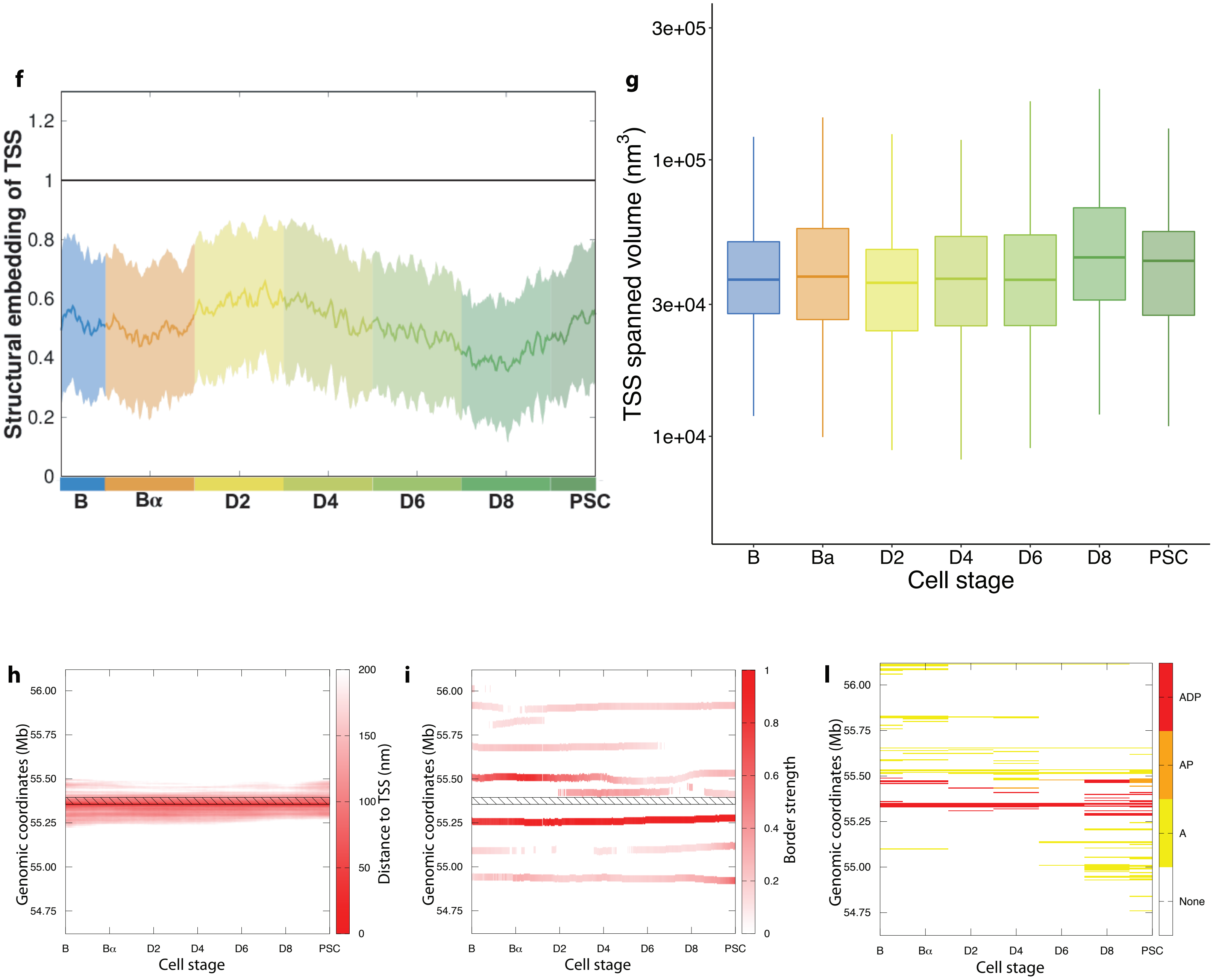


**m**

| TSS | B | Bα | D2 | D4 | D6 | D8 | PSC |
|---|---|---|---|---|---|---|---|
| **A** | 31 | 26 | 16 | 23 | 13 | 36 | 47 |
| **AP** | 10 | 8 | 7 | 8 | 5 | 14 | 18 |
| **APD** | 10 | 8 | 7 | 7 | 4 | 14 | 12 |

### Supplemental figure 10

##### Sox2 chr3:34649995-34652460. Forward

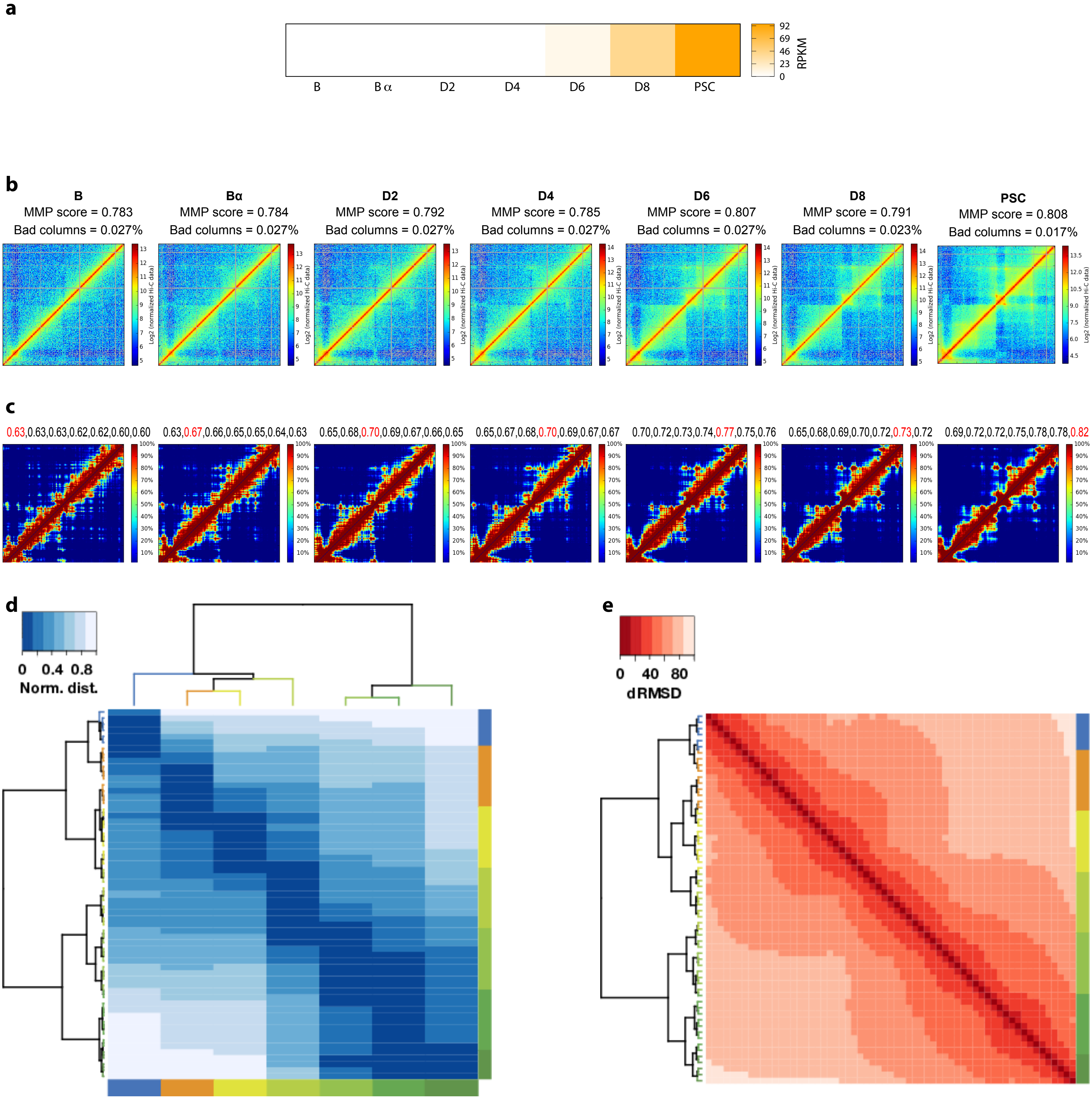

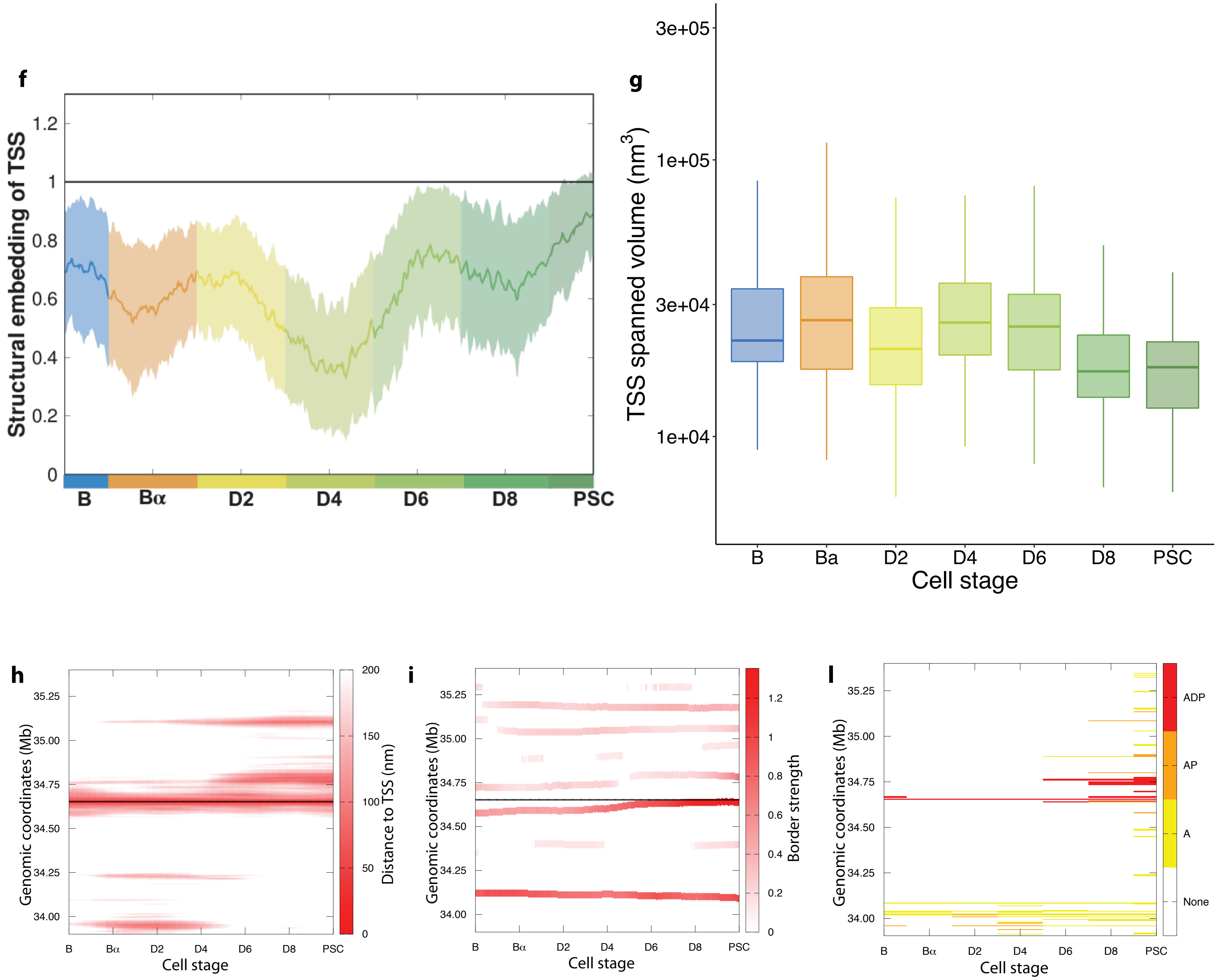


**m**

| TSS | B | Bα | D2 | D4 | D6 | D8 | PSC |
|---|---|---|---|---|---|---|---|
| **A** | 9 | 6 | 7 | 13 | 13 | 22 | 48 |
| **AP** | 4 | 1 | 4 | 4 | 4 | 13 | 23 |
| **APD** | 3 | 1 | 1 | 1 | 4 | 10 | 15 |

### Supplemental figure 11

#### Tet2 chr3:133463679-133545139. Reverse

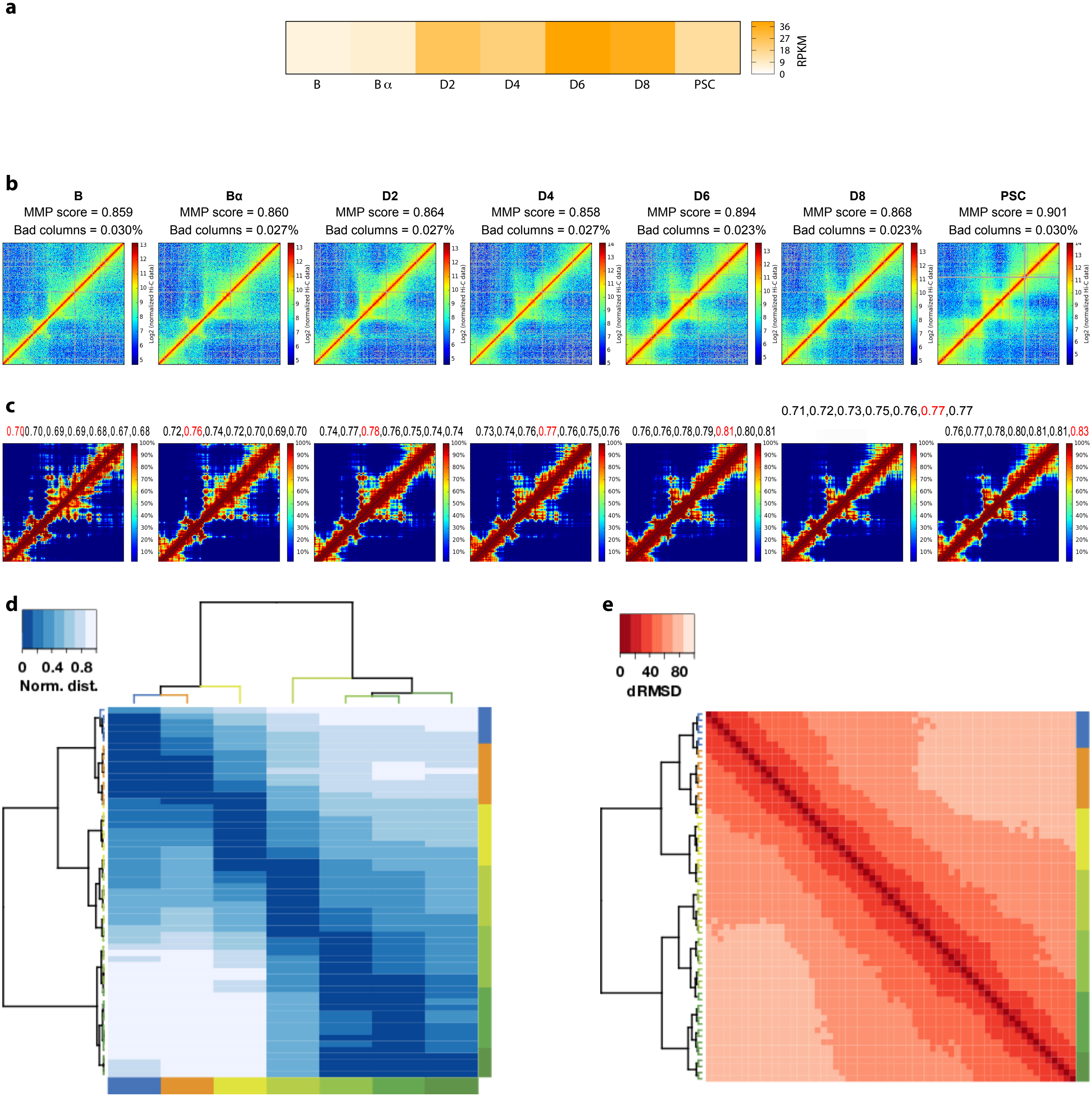

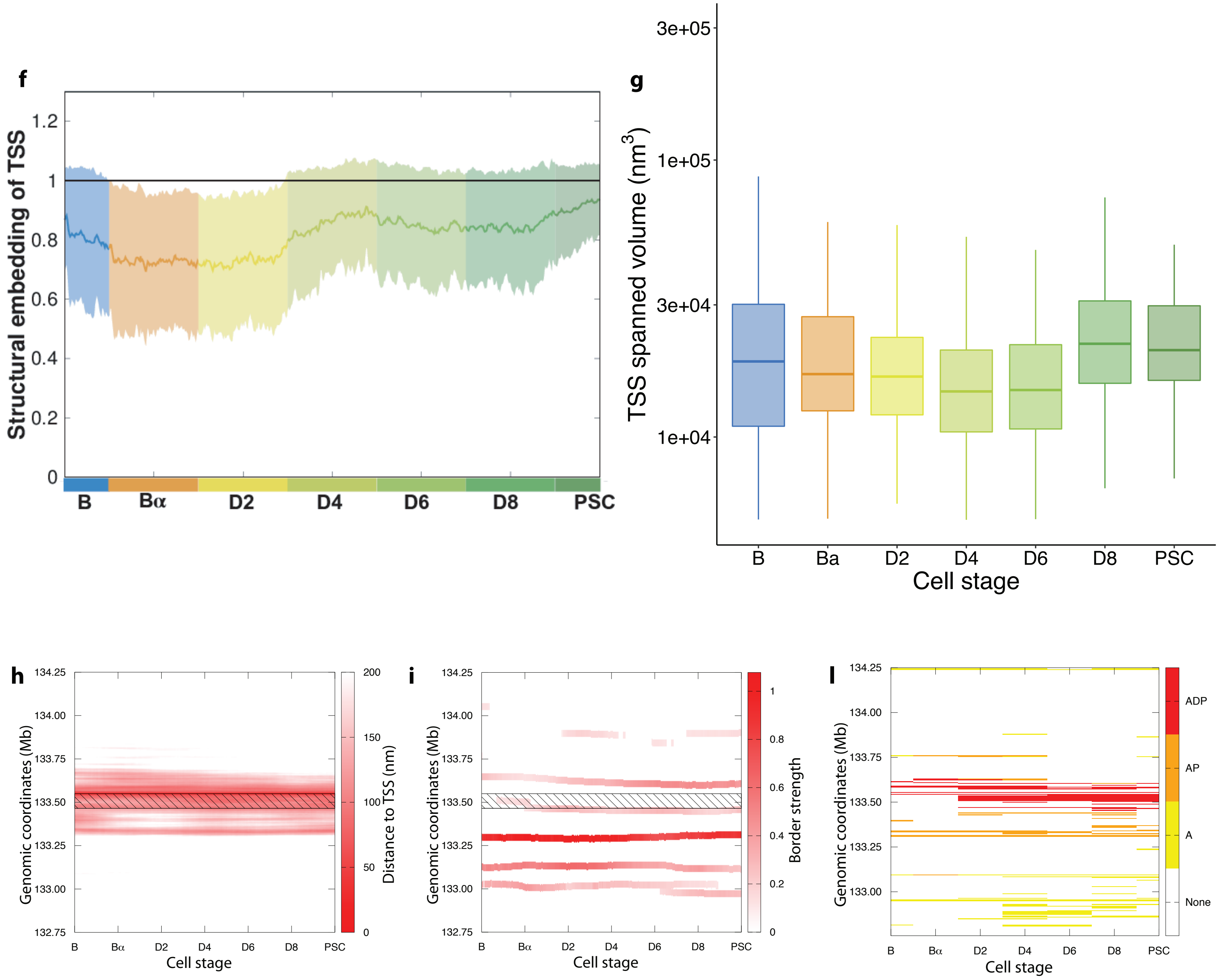


**m**

| TSS | B | Bα | D2 | D4 | D6 | D8 | PSC |
|---|---|---|---|---|---|---|---|
| **A** | 18 | 17 | 31 | 47 | 32 | 45 | 37 |
| **AP** | 11 | 13 | 24 | 25 | 18 | 29 | 21 |
| **APD** | 5 | 7 | 15 | 15 | 13 | 18 | 15 |
